# The RNA helicase HrpA rescues collided ribosomes in *E. coli*

**DOI:** 10.1101/2024.09.11.612461

**Authors:** Annabelle Campbell, Hanna F. Esser, A. Maxwell Burroughs, Otto Berninghausen, L. Aravind, Thomas Becker, Rachel Green, Roland Beckmann, Allen R. Buskirk

## Abstract

Although many antibiotics inhibit bacterial ribosomes, loss of known factors that rescue stalled ribosomes does not lead to robust antibiotic sensitivity in *E. coli*, suggesting the existence of additional mechanisms. Here, we show that the RNA helicase HrpA rescues stalled ribosomes in *E. coli.* Acting selectively on ribosomes that have collided, HrpA uses ATP hydrolysis to split stalled ribosomes into subunits. Cryo-EM structures reveal how HrpA simultaneously binds to two collided ribosomes, explaining its selectivity, and how its helicase module engages downstream mRNA, such that by exerting a pulling force on the mRNA, it would destabilize the stalled ribosome. These studies show that ribosome splitting is a conserved mechanism that allows proteobacteria to tolerate ribosome-targeting antibiotics.

## Introduction

Translating ribosomes often stall during protein synthesis in bacteria. Although some pauses are reversible, and even desirable, promoting protein folding^1^ and regulating gene expression,^2–4^ prolonged stalling on truncated or damaged mRNAs traps ribosomes in an inactive state and produces truncated, potentially toxic peptides.^5^ Ribosome rescue pathways selectively recognize arrested ribosomes, recycle the ribosome subunits, and target the aborted nascent peptide for degradation.^6^

We recently showed that in bacteria, like eukaryotes,^7,8^ ribosome collisions trigger ribosome rescue pathways.^9,10^ In bacteria, collisions are recognized by SMR-domain proteins that have two different domain architectures. The simple SmrB architecture, with only an SMR domain and N-terminal extension, is mainly found in proteobacteria (Fig. S1A). In *E. coli*, for example, SmrB is recruited to collided ribosomes, where its SMR domain cleaves the mRNA, targeting it for decay.^9^ When upstream ribosomes reach the newly formed 3’-end of the mRNA, they are rescued by tmRNA, which tags the nascent peptide for degradation and recycles the ribosomal subunits. In contrast, the more complex MutS2 architecture is found in many other bacterial phyla (Fig. S1A). Our recent work in *B. subtilis* revealed that the SMR domain of MutS2 functions only as a collision sensor, not as a nuclease.^10^ Instead, MutS2 uses its ABC ATPase domain to split the leading ribosome into subunits.^10,11^ Peptidyl-tRNA trapped on the large ribosomal subunit is then recognized by RqcH and RqcP, which add Ala residues to the end of the nascent peptide, targeting it for degradation.^12–14^

One potential advantage of ribosome splitting over mRNA cleavage and tmRNA-mediated rescue is that it does not require the ribosome to be active. This is likely relevant in cells treated with antibiotics that target elongating ribosomes. Indeed, *B. subtilis* cells lacking MutS2 are hypersensitive to such antibiotics,^10,11^ whereas *E. coli* cells lacking SmrB are not. These observations suggest that *E. coli* has an unknown mechanism essential for rescuing antibiotic-arrested ribosomes. Here we show that this mechanism involves a new ribosome rescue factor in *E. coli*, HrpA, a DExH-box ATPase with 3’-5’ RNA helicase activity.^15^ Genetic screens suggested that HrpA may play a role in protein synthesis^16,17^ and in tolerance to antibiotics that target ribosomes.^18,19^ Here we define the molecular mechanism for HrpA in ribosome rescue. We show that HrpA rescues ribosomes stalled by arrest peptides on reporter mRNAs as well as ribosomes stalled transcriptome-wide by antibiotics. HrpA splits ribosomes into subunits *in vitro* and this activity is dependent on its unique C-terminal region and its ability to bind and hydrolyze ATP. We show that this activity is specific for collided ribosomes. Finally, we solve the cryo-EM structure of HrpA bound to collided ribosomes, providing insight into its selectivity and its molecular mechanism.

## Results

### Δ*hrpA* cells are hypersensitive to antibiotics that target ribosomes

To confirm previous reports of antibiotic sensitivity in cells lacking HrpA,^15,18^ we plated cells from strains derived from *E. coli* K12 MG1655 on media containing antibiotics that target the elongation stage of protein synthesis by different mechanisms. Although the wild-type strain survives robustly at low concentrations of chloramphenicol or erythromycin, the Δ*hrpA* strain shows little or no growth (Fig. 1A). The Δ*hrpA* strain is also hypersensitive to fusidic acid, and its growth is impaired significantly, if not entirely, by tetracycline and spectinomycin. In contrast, for each of the antibiotics tested, the Δ*smrB* strain grows about as well as the wild-type strain (Fig. 1A), arguing that SmrB’s RNase activity is not essential for cells to respond to ribosome stalling induced by antibiotics. These results with a variety of elongation inhibitors suggest that HrpA and SmrB work in different pathways or by different mechanisms, and that HrpA plays a more important physiological role in response to antibiotic treatment.

**Fig. 1.**
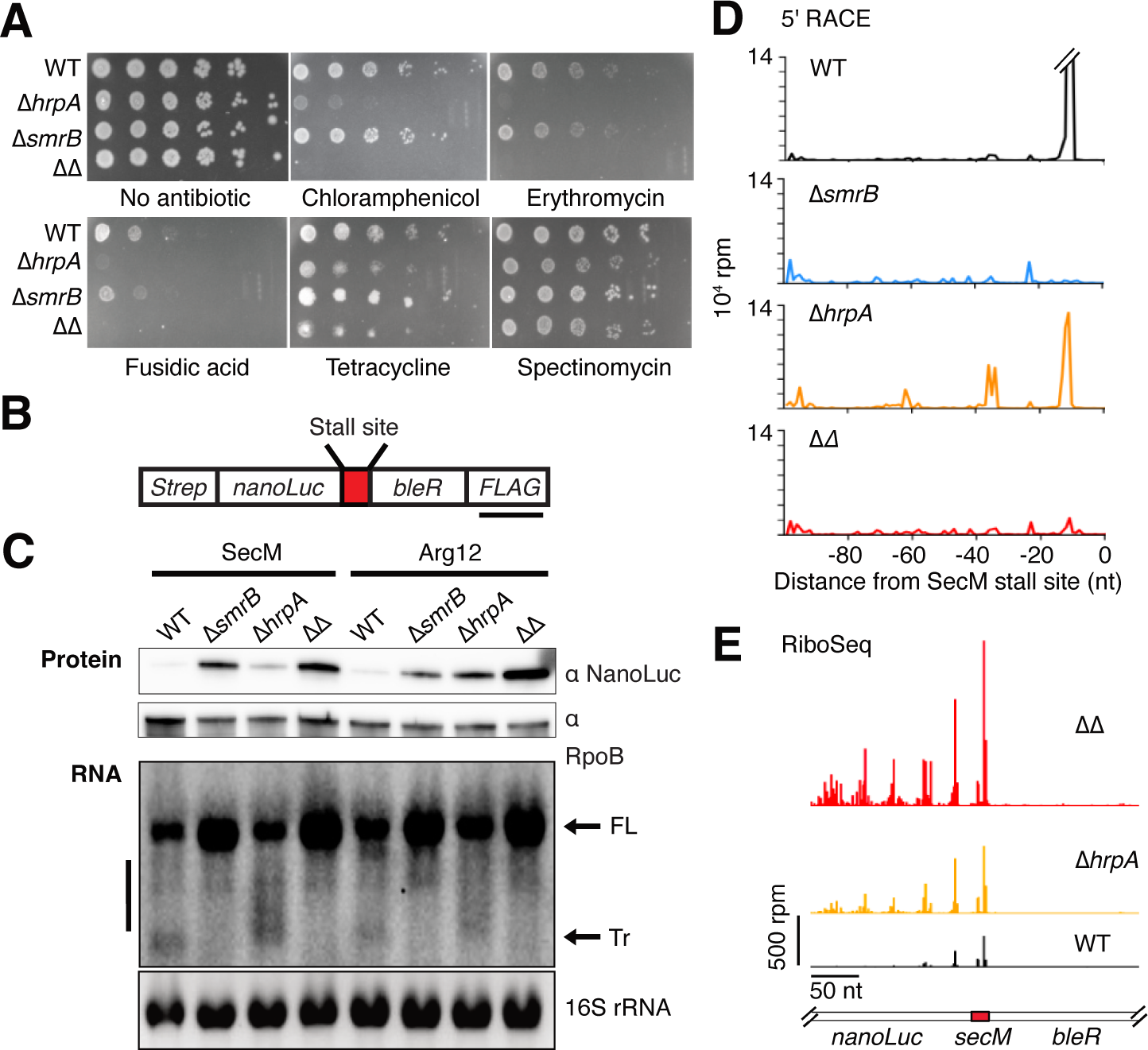
HrpA removes stalled ribosomes from reporter mRNAs. (**A**) Growth assays of cells on chloramphenicol (2.5 µg/mL), erythromycin (100 µg/mL), fusidic acid (100 µg/mL), tetracycline (0.8 µg/mL), and spectinomycin (12.5 µg/mL). (**B**) Reporter expressing a fusion of NanoLuc and bleomycin resistance protein with a stalling motif (SecM or Arg12) inserted between them. (**C**) Top: Immunoblot showing full-length reporter protein levels detected with anti-NanoLuc antibody. The loading control was RpoB. Bottom: Northern blot showing reporter mRNA levels detected with a 3’ probe. The loading control was 16S rRNA. FL = Full-length. Tr = truncated. The vertical bar spans longer mRNA fragments. (**D**) 5ʹ-RACE data showing the 5ʹ ends of downstream fragments in reads per million (RPM) on the SecM reporter. The first nucleotide in the A-site codon in the stalling motif was designated position 0. (**E**), Ribosome profiling data showing ribosome density in RPM on the SecM reporter.

### HrpA removes stalled ribosomes on reporter mRNAs

We next asked how HrpA affects the expression of reporter constructs in which ribosomes are arrested at specific sites. We employed a set of mRNA reporters containing the nano-luciferase gene fused to the bleomycin resistance gene in the same open reading frame.^9^ Between them we inserted the SecM arrest peptide or a tract of twelve rare arginine codons (Arg12) to arrest translating ribosomes (Fig. 1B). The SecM and Arg12 sequences are known to robustly arrest the ribosome, consistent with the low protein output from these reporters.^2,9,20^ In immunoblots using antibodies against NanoLuc, we observe a strong increase in the protein levels from both reporter constructs in the Δ*smrB* strain, as reported previously^9^ (Fig. 1C, top). Similarly, Δ*hrpA* cells show a small increase in the amount of protein from the SecM reporter and a larger increase with the Arg12 reporter. Finally, cells lacking both factors (ΔΔ) show an additive increase in protein from both reporters.

To determine the effects of SmrB and HrpA on the reporter mRNAs, we performed northern blots using probes complementary to the 3’-end of the reporter transcripts (Fig. 1C, bottom). In the Δ*smrB* strain and the double knockout (ΔΔ), the full-length reporter mRNA is stabilized and the truncated fragment downstream of the stall site is no longer observed. We previously showed that SmrB cleaves mRNA between collided ribosomes, promoting mRNA decay.^9^ The higher level of protein output in the Δ*smrB* strains is likely due to these higher levels of reporter mRNA. In contrast, there is no change in the level of full-length mRNA in the Δ*hrpA* strain, arguing that its mechanism of action is not primarily on mRNA decay.

In the Δ*hrpA* strain, in addition to full-length mRNA, we observed a broad distribution of mRNA fragments longer than the truncated band in the wild-type strain (Fig. 1C, bottom). These fragments disappear when both *smrB* and *hrpA* are deleted, suggesting that SmrB cleaves the reporter mRNA differently in the Δ*hrpA* strain. To determine where this cleavage occurs, we used 5’-rapid amplification of cDNA ends (RACE) on the SecM reporter in the various strains. As expected based on our previous biochemical and structural studies,^9^ the 5’-end of the downstream fragment produced by SmrB is 11 nt upstream of the SecM stall site, corresponding to the 5’-boundary of the stalled ribosome (Fig. 1D). In the absence of HrpA, there are additional sequencing reads (5’-ends) at ∼25 nt intervals upstream of the first peak. Given that a bacterial ribosome footprint is about 25 nt,^21^ we conclude that there is a longer queue of collided ribosomes upstream of the stall site in the absence of HrpA, and that SmrB cleaves the mRNA between them. Consistent with this model, all peaks are lost in the Δ*hrpA* Δ*smrB* strain.

Two additional experiments suggest that stalled and collided ribosomes accumulate in cells lacking HrpA. First, ribosome profiling data from strains expressing the SecM reporter show an increase in ribosome density upstream of the stall site in the absence of HrpA, consistent with four or five ribosomes stacking behind the SecM-stalled ribosome (Fig. 1E). These effects are even clearer in the Δ*hrpA* Δ*smrB* strain, due to the increased number of reads from the reporter mRNA. Second, we detected SmrB across sucrose gradients from cells treated with different concentrations of erythromycin (Fig. S2). In each condition, the collision-binding protein SmrB migrates deeper in the gradients in the Δ*hrpA* strain, suggesting that there are more ribosome collisions in the absence of HrpA. Taken together, these data are consistent with HrpA removing stalled ribosomes from mRNAs and aborting protein synthesis, whether the ribosomes are stalled by arrest peptides or rare codons on the reporter mRNAs or by antibiotics across the transcriptome.

### HrpA selectively rescues collided ribosomes

How does HrpA distinguish stalled ribosomes from actively elongating ribosomes? To test the hypothesis that, like SmrB, HrpA selectively recognizes collided ribosomes, we generated reporters with varied distances from the start site to the Arg12 stalling motif (39, 99, or 210 nt, Fig. 2A). The closer the stalling motif is to the start codon, the less room there is for ribosomes to accumulate and collide.^7,9^ In immunoblots using antibodies against the C-terminal FLAG epitope on the reporter protein, we observed significantly more full-length protein in the Arg12-99 and Arg12-210 reporters in the Δ*hrpA* strain than in the wild-type strain (Fig. 2B); we interpret these data to mean that without HrpA, stalled ribosomes are not removed from the mRNA, and they eventually read through the Arg12 stalling sequence. In contrast, with the Arg12-39 reporter, where at most two ribosomes can be loaded on the stall site, there are similar amounts of protein produced in the Δ*hrpA* and wild-type strains, suggesting that HrpA cannot rescue ribosomes in the absence of collisions. Similarly, loss of SmrB only slightly increased protein output from the Arg12-39 and Arg12-99 reporters, but dramatically increased the output from the Arg12-210 reporter (Fig. 2B). The observation that loss of HrpA has a greater effect on the Arg12-99 reporter than the loss of SmrB suggests that HrpA may resolve collisions containing queues of fewer ribosomes than SmrB requires. Finally, for the Arg12-210 reporter, seemingly a good substrate for both factors, the double knockout shows an additive effect, underscoring that these two factors work by independent mechanisms.

**Fig. 2.**
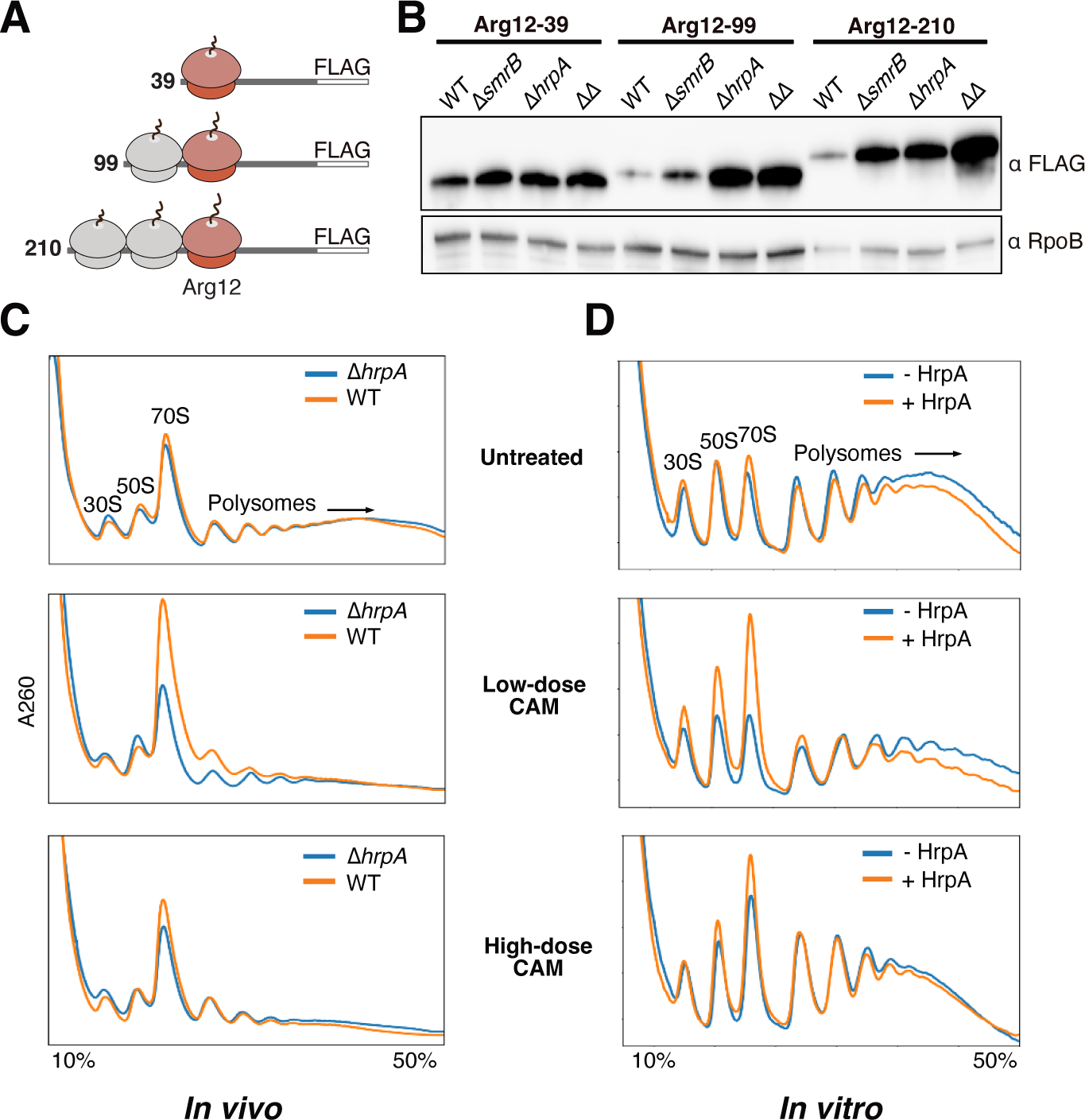
HrpA splits collided ribosomes. (**A**) Reporters with varied distance from the start site to the Arg12 stalling sequence. (The longest constructs should accommodate more than three ribosomes). (**B**) Immunoblot showing the full-length reporter protein detected with anti-FLAG antibody. The loading control was RpoB. (**C**) Polysome traces from untreated, low-dose (5 µg/ml), or high-dose (500 µg/ml) chloramphenicol (CAM)-treated cells. (**D**) Polysome traces from *in vitro* reactions with 0 or 0.5 pmol HrpA and 25 pmol ribosomes from polysomes from untreated, low-dose, or high-dose CAM-treated cells.

To gain a broader view of HrpA across the transcriptome, we used an antibiotic titration to distinguish the effects of ribosome stalling or ribosome collisions. When cells are treated with a low dose of an elongation inhibitor, some ribosomes stall, and uninhibited ribosomes collide with them; by contrast, in cells treated with high doses of inhibitor, few ribosomes collide because all ribosomes are quickly inhibited.^7,9^ For this experiment, lysates from cultures treated with chloramphenicol (CAM) were analyzed on sucrose gradients. We did not observe a marked difference between the absorbance signal from polysome profiles of wild-type and Δ*hrpA* cells in untreated conditions or at the high dose (Fig. 2C). In contrast, low doses of CAM decreased deep polysomes while increasing monosomes, disomes, and trisomes in the wild-type strain, but not the Δ*hrpA* strain (Fig. 2C, middle). These results argue that HrpA affects translation in response to ribosome collisions induced by antibiotics.

We hypothesize that HrpA splits stalled ribosomes into their component subunits. If HrpA splits one ribosome in a collided pair, we expect to see a decreased polysome signal and a corresponding increase in both ribosome subunits and monosomes. Interestingly, we only see an increase in monosomes in Fig. 2C (middle), perhaps because subunits reassociate into empty 70S complexes in cell lysates. To better characterize the products of HrpA activity, we performed a similar experiment *in vitro* using ribosomes pelleted from Δ*hrpA* lysates under conditions that enrich for polysomes with or without collisions. These isolated polysomes were incubated with purified HrpA at approximately physiological levels relative to ribosomes (∼1:50), ATP, and an excess of IF3 to prevent subunit reassociation. The reactions were then analyzed on sucrose gradients. As observed *in vivo*, HrpA has little effect on ribosomes from untreated or high-dose CAM treated lysates (Fig. 2D). Strikingly, however, incubation of HrpA with polysomes from low-dose CAM-treated cells led to decreased polysomes and increased abundance of both subunits and monosomes (Fig. 2D, middle). These data provide strong support for our hypothesis that HrpA splits collided ribosomes.

### Structure of HrpA-bound collided disomes

To gain deeper mechanistic insight into how HrpA specifically recognizes and splits collided ribosomes, we translated an mRNA encoding the VemP arrest peptide^9,22^ in an *in vitro* reconstituted *E. coli* translation system (PURE) and isolated the disome peaks from sucrose gradients. As expected, collided ribosomes prepared in the PURE system are split into subunits in the presence of HrpA and ATP (Fig. S3). To trap the complex prior to splitting, HrpA and the non-hydrolysable ATP analogue AMPPNP were added to the prepared VemP collided ribosomes, and the sample was subjected to analysis by cryo-EM. Analysis of single particles revealed disomes with tRNAs in the classical state (A/A and P/P) in the stalled ribosome, as observed previously,^9,23^ with the collided ribosome in either the classical or hybrid (A/P and P/E tRNA) state (Fig. S4). Additional density was observed bridging the stalled and collided ribosomes, corresponding to one or two copies of HrpA (Fig. S4). In class I, with one copy of HrpA, the disome structure was refined to an overall resolution of 3.1 Å with local resolution for HrpA ranging from 4-18 Å (Fig. 3A, Fig. S5). This resolution allowed us to fit an AlphaFold2 model^24^ as well as a crystal structure^15^ of HrpA into the final map (Fig. 3A, Figs. S6 and S7, table S1). In class II (3.2 Å), a second HrpA molecule was partly visible as described below (Figs. S4, S5, S6F and S8).

**Fig. 3.**
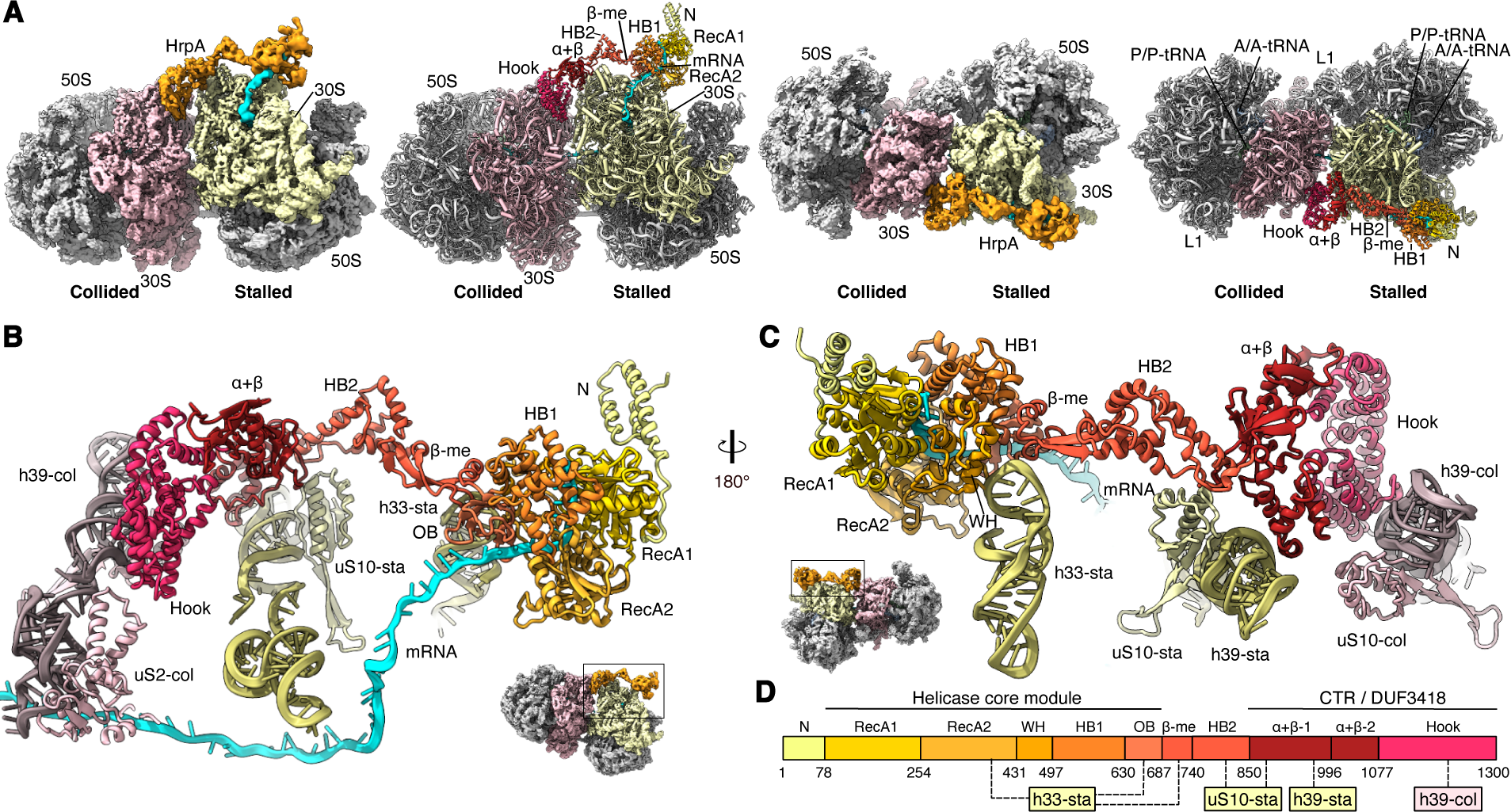
Structure of the HrpA-bound disome complex. (**A**) Cryo-EM density map and molecular model of the *E. coli* HrpA-bound disome shown as side (left panels) or top views (right panels). The composite map consists of the refined disome, gaussian low-pass filtered isolated maps for HrpA Hook/α+β and β-meander/HB2 domains as well as a focused refined map for the HrpA helicase domain (see Figs. S4-7). Isolated densities for HrpA are shown at lower contour levels compared to the disome for clarity. A color code for HrpA domains is given in **D**. (**B** and **C**) Molecular model demonstrating the recognition of the stalled disome by HrpA. The model is shown as a front view (**B**) and a back view (**C**) as indicated in the thumbnails. Herein, the black box shows the zoomed region. (**D**) Schematic representation of the HrpA domain organisation and interactions with the disome. sta, stalled 70S; col, collided 70S.

In both structures, HrpA stretches from the mRNA entry site of the stalled ribosome across the 30S head until it reaches the 30S subunit of the collided ribosome, a distance of about 150 Å (Fig. 3A, Fig. S8). The core module of HrpA, shared with a broad range of RNA helicases, contains the following domains: a N-terminal 3-helix bundle (N, residues 1-78), the active P-loop NTPase with an intact Walker A and B motif (RecA1, 79-254), an inactive P-loop NTPase (RecA2, 255-431), a winged helix-turn-helix (wHTH, 432-497), a helical bundle formed by a helix-hairpin-helix and HTH domain (HB1, 499-630), and an OB-fold (631-687). This core module is positioned right above the mRNA entry site, on the beak of the 30S head of the stalled ribosome. The RecA2 and OB-fold domains interact with rRNA in helix h33, as does the β-meander domain (residues 688-740, Fig. 3C, D). Strikingly, from the mRNA entry site we observe mRNA density that leads directly into the helicase core of HrpA (Fig. 3, Fig. S7). Comparisons with structures of HrpA in the RNA-bound conformation (PDB-ID 6zww)^15^ and the unbound conformation (PDB-ID 6zwx)^15^ further establish that the two RecA domains in the helicase module are engaged with mRNA in our structure (Fig. S7). We conclude that HrpA engages the mRNA protruding from the stalled ribosome at the mRNA entry site.

HrpA recognizes ribosome collisions by binding both the stalled and collided ribosomes via its unique C-terminal region (CTR). This region belongs to the DUF3418-like superfamily^25^ and features the following three domains: α+β-1 which resembles an inactive restriction endonuclease domain (851-996), α+β-2 (997-1077), and a helical domain we refer to as the Hook domain (1078-1300). The Hook domain is positioned at the disome interface between the 30S heads (Fig. 3A-C). The main contact of HrpA with the collided ribosome is formed by the C-terminal 3-helix-element of the Hook with the back of the 30S head, where it interacts with 16S rRNA helix h39 located between r-proteins uS10 and uS2 (Fig. 3B-D). Additional contacts between HrpA and the stalled ribosome are formed by the two α+β domains and HB2 with h39 and uS10 in the stalled ribosome (Fig. 3C, D). The CTR is connected to the N-terminal helicase core module via a second helical bundle (HB2, 741-850) and β-meander (residues 688-740) domains.

Class II contains one molecule of HrpA in an essentially identical conformation as in Class I. In addition, extra density was present for parts of a second HrpA (HrpA2, Figs. S6F, G and S8), corresponding to the Hook domain and parts of the second α+β domain. Hook2 packs against both the stalled and collided ribosomes utilizing a different surface than Hook1 (Fig. S8E, F). Although the N-terminal domains of HrpA2 are largely unstructured, we observe extra density adjacent to the HB2, β-meander, and OB-fold domains of HrpA1 reaching towards the mRNA entry channel.

### Requirements for HrpA to split ribosomes into subunits

To characterize the contributions of various regions of HrpA to its catalytic activity, we purified and characterized a series of HrpA mutants. We increased the signal in our *in vitro* splitting assay by raising the amount of HrpA to levels approximately equal to the ribosomes in the collision-enriched polysomes. With this increased amount of HrpA, there is a striking decrease in polysomes and an increase in monosomes and subunits (Fig. 4A). As expected, the splitting reaction depends on the presence of ATP (Fig. 4A). We tested mutations in the first RecA domain designed to disrupt ATP binding (Walker A, K106A) or hydrolysis (Walker B, D197A). The K106A and D197A mutants both show little or no splitting activity, arguing that both ATP binding and hydrolysis are required for HrpA activity (Fig. 4B).

**Fig. 4.**
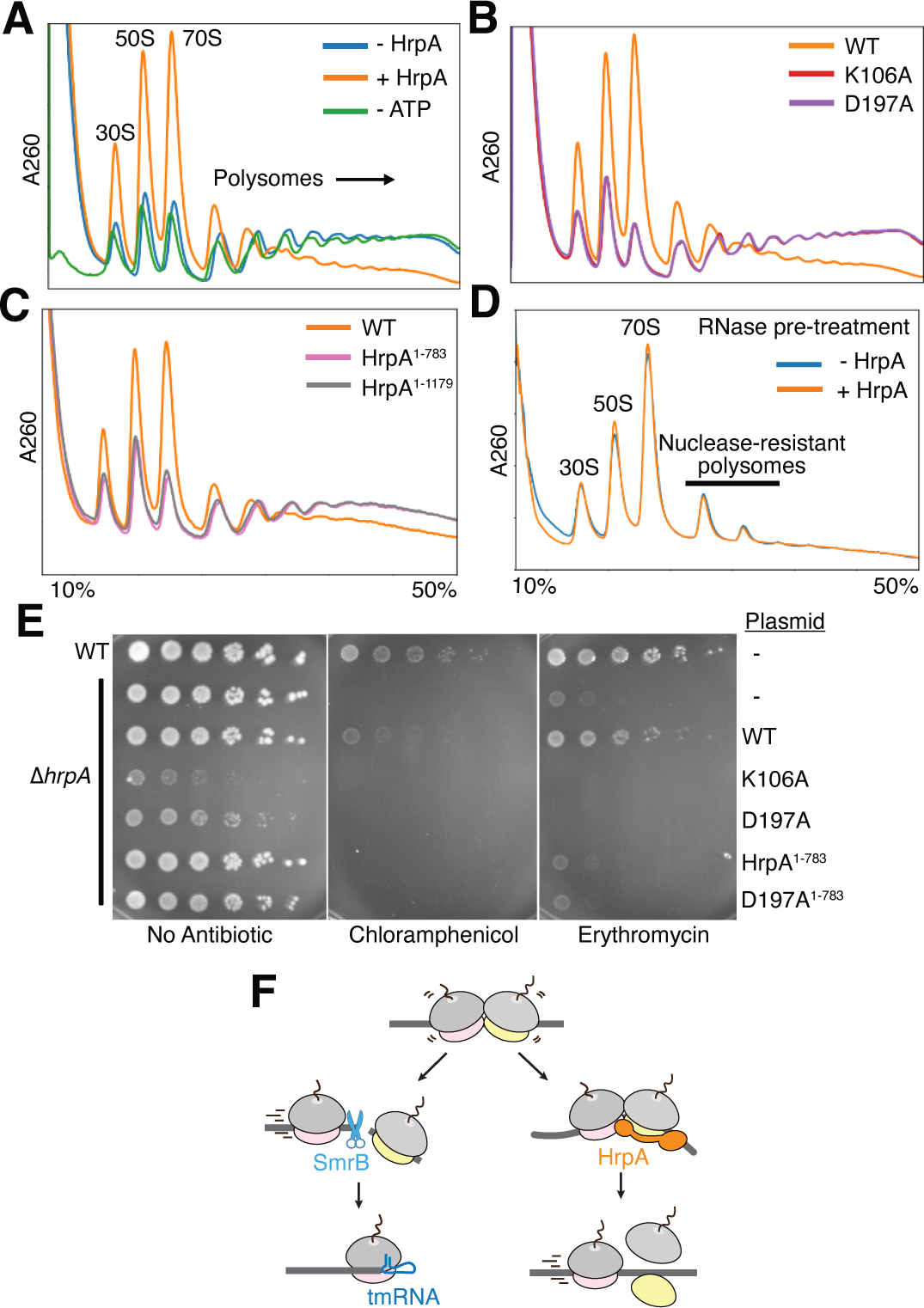
HrpA requires ATP, its CTR, and mRNA for splitting. Polysome traces from *in vitro* reactions using 25 pmol polysomes enriched from low-dose (5 µg/ml) CAM-treated cells. All reactions contain 250 pmol IF3 and 1 mM ATP unless otherwise indicated. (A) Reactions with 0 or 25 pmol wild-type HrpA. The - ATP reaction contained 25 pmol wild-type HrpA and no ATP. (**B** and **C**) Reactions contained 25 pmol wild-type or mutant HrpA. (**D**) Polysomes were pre-treated with RNase A before incubation with 0 or 2.5 pmol HrpA. (**E**) Growth assays on chloramphenicol (2.5 µg/mL) and erythromycin (100 µg/mL). (**F**) Model for the two ribosome rescue pathways in *E. coli*.

We next looked at HrpA-ribosome contact sites revealed by the cryo-EM structure. To test whether the interaction between the CTR and the collided ribosome is required for binding, we expressed FLAG-tagged HrpA constructs in Δ*hrpA* cells (Fig. S9A) and observed their distribution on sucrose gradients. Wild-type HrpA is present in nearly all fractions from the sucrose gradient (Fig. S9B). In contrast, less of the HrpA^1–783^ mutant lacking the CTR is present in polysome fractions, suggesting it does not bind ribosomes as well. We performed the same experiment with an HrpA mutant lacking the N-terminal domains (HrpA^784–1300^) and saw similar results. These data suggest that both regions are required for optimal binding (Fig. S9B).

We also tested whether C-terminal deletions impact HrpA’s splitting activity. We found that the HrpA^1–783^ mutant (depicted in Fig. S6h) cannot split collided ribosomes *in vitro* (Fig. 4C) or *in vivo* (Fig. S9C). These observations are consistent with the binding defects observed above and with an earlier report that expression of the HrpA^1–783^ mutant does not complement the Δ*hrpA* strain even though it maintains full helicase activity *in vitro.*^15^ More specifically, our structure revealed that the C-terminal helical bundle in the Hook domain contacts h39 of the collided ribosome. We purified an HrpA mutant lacking the C-terminal 121 amino acids of the Hook domain (Fig. S6H) and found that this HrpA^1–1179^ mutant is also unable to split ribosomes *in vitro*, suggesting the helical bundle region is important for detecting collided ribosomes (Fig. 4C).

The structure also revealed that the helicase region of HrpA engages mRNA downstream of the stalled ribosome. Given that HrpA is a 3’ to 5’ helicase, we wondered if it requires mRNA downstream of ribosome collisions to grasp and destabilize the complexes to catalyze subunit splitting. We tested this with our *in vitro* assay. Unlike reactions with intact mRNA, where splitting yields high levels of monosomes and subunits, when we add RNase A to polysomes prior to incubation with HrpA, ATP, and IF3, there is no increase in monosomes or subunits compared to the minus HrpA control (Fig. 4D). Moreover, there is no change in the level of nuclease-resistant disomes and trisomes, suggesting that an mRNA handle is required for HrpA to split collided ribosomes (Fig. 4D).

Finally, we analyzed the antibiotic sensitivity of cells expressing HrpA mutants (Fig. 4E). Ectopic expression of wild-type HrpA in Δ*hrpA* cells rescues the hypersensitivity to chloramphenicol and erythromycin. Expression of the K106A or the D197A mutants did not complement the deletion strain; rather, it confers a dominant negative phenotype (on plates with no antibiotic, Fig. 4D) and disrupts general translation, yielding high levels of separate subunits in sucrose gradients (Fig. S9D). As expected, the CTR truncation mutant, HrpA^1–783^, does not rescue the deletion phenotype, consistent with its lack of splitting activity in vitro. Interestingly, deleting the C-terminal region of the Walker B mutant (D197A^1–783^) relieves the dominant negative toxicity observed in the D197A mutant. These in vivo results broadly validate the structural observations and in vitro biochemistry with the mutants, underscoring the importance of ATP binding and hydrolysis and the C-terminal region for HrpA activity.

## Discussion

Here we show that HrpA is a ribosome rescue factor that splits stalled 70S ribosomes into subunits (Fig. 4F). In eukaryotes, stalled 80S ribosomes are split by a distinct helicase known as Slh1^26–29^ in a striking example of functional convergence. HrpA is not the bacterial ortholog of Slh1; it is more closely related to the eukaryotic Dhx/Dhr/Prp helicases, sharing the core N-terminal domains detailed above.^30^ Acquisition of diverse C-terminal domains across this clade led to their functional specialization in diverse pathways. In eukaryotes, the C-terminal domains recruited the Dhx/Dhr/Prp helicases to ribosome biogenesis and splicing pathways. In bacteria, the CTR domains recognizing the ribosome collision interface likely recruited HrpA to the ribosome rescue pathway. In contrast, Slh1 does not sense collided ribosomes directly; it is part of the RQT complex that binds ubiquitin chains added by the E3-ligase Hel2.^26,31^

To promote the splitting reaction, HrpA’s helicase module is anchored to the ribosome near the beak and engages downstream mRNA such that its 3’ to 5’ helicase activity^15^ can exert a pulling force on the mRNA out of the ribosome (Fig. S10). This generation of force by the helicase would likely rotate the head of the 30S subunit towards the 50S subunit, a conformational change that could destabilize the interface of the stalled ribosome. In a similar manner, the RQT complex binds to the ribosome near the mRNA entry site (Fig. S10) and requires protruding mRNA to engage.^28^ Although no engaged mRNA has been directly observed in the eukaryotic system, as it was here in the bacterial system, an inward rotation of the 40S head was observed in cryo-EM analyses of splitting reactions.^28^ While HrpA has a single helicase cassette, Slh1 contains two helicase cassettes and Walker mutations in either one renders the complex inactive.^28^ Further studies will be required to determine the functional consequences of the second HrpA molecule seen in a sub-population of our particles.

Ribosome splitting is the dominant pathway for ribosome rescue in yeast and animals^26,27,32^ and HrpA may be the first line of defense in *E. coli* as well. Broadly speaking, HrpA is encoded in most proteobacterial genomes that encode SmrB. Perhaps because it can bind between each ribosome pair, SmrB binds and cleaves more efficiently when there are three or more ribosomes.^9^ In contrast, HrpA will only act on the first stalled ribosome, no matter how long the queue, because it must engage the downstream mRNA which is not accessible between collided ribosomes. Thus, mRNA cleavage may play a more critical role when long ribosome queues form due to irreversible arrest or mRNA damage and where HrpA is overwhelmed.

Which substrates are the most relevant for the various bacterial ribosome rescue factors? We envision that HrpA rescues collided disomes arrested by any failure in translation where ribosomes stall on the mRNA, but that it may play a more important role for some stalled complexes than others. Ribosome splitting leaves the mRNA intact and gives upstream ribosomes a chance to complete protein synthesis. This solution seems appropriate for antibiotic-stalled ribosomes, especially given that subunit splitting does not require ribosome activity. In contrast, following SmrB cleavage, upstream ribosomes on the truncated mRNA are rescued by tmRNA, which requires ongoing, active translation, and thus cannot happen with antibiotic-stalled ribosomes. These considerations may explain the striking difference in antibiotic sensitivities of strains lacking SmrB or HrpA. Additionally, mRNA cleavage by SmrB may be more critical for damaged mRNAs or situations where HrpA cannot engage, such as for collided ribosomes stalled behind RNA polymerase in coupled transcription-translation complexes.^33^ Future work is needed to clarify the distinct contributions of SmrB and HrpA to translational homeostasis.

## Acknowledgments

The authors thank the JHMI Single Cell and Transcriptomics for assistance with high-throughput sequencing and A. Mao and M. Catipovic (JHU) as well as C. Ungewickell, S. Rieder, and Joanna Musial (LMU) for excellent technical assistance. This work was supported by NIH grants F31AI172391 (A.C.) and R01GM136960 (A.R.B.), HHMI (R.G.), the German Research Council (BE1814-20-1 and BE1814-22-1, R.B.), the ERC (ADG 885711, R.B.), and the Intramural Research Program of the National Library of Medicine at the NIH (A.M.B. and L.A.).

## Author contributions

A.C. identified HrpA as a potential ribosome rescue factor and performed the antibiotic sensitivity assays, the analysis of the stalling reporters, and the subunit splitting assays in vitro. A.M.B. and L.A. performed the phylogenetic analyses. H.F.E. prepared samples for cryo-EM analysis and processed the cryo-EM data. O.B. collected the cryo-EM data. H.F.E. prepared the molecular models. H.F.E., T.B., and R.B. analyzed and interpreted the structures, and H.F.E. and T.B. prepared the structural figures. R.G., R.B., and A.R.B. supervised the project and with A.C, H.F.E, and T.B. wrote the paper.

## Supplemental Information

Figures S1 – S10

Table S1

## Materials and methods

### Data availability

The cryo-EM structural data generated in this study have been deposited in the Protein Data Bank and the Electron Microscopy Data Bank under accession codes EMD-XXXXX and PDB-XXXX for the HrpA-disome complex as well as EMD-YYYYY and PDB-YYYY for the (HrpA)2-disome complex. The ribo-seq data were deposited in the NCBI Gene Expression Omnibus database with accession code GSE270401.

### Code availability

Custom python scripts used to analyse the ribo-seq data are freely available at: https://github.com/greenlabjhmi/2024_HrpA.

### Statistics and reproducibility

Western blots (Figs. 1 & 2), northern blots (Fig. 1), and polysome profiles (Figs. 2 & 4) were repeated at least three times. The 5’ RACE and ribosome profiling experiments were performed in duplicate from biological replicates (Fig. 1). The in vitro HrpA splitting assays (Figs. 2 & 4) were performed in triplicate. All attempts at replication were successful.

### Sequence Analyses

PSI-BLAST (RRID:<u>SCR_001010</u>)^34^ and JACKHMMER (RRID:<u>SCR_005305</u>)^35^ were used to carry out iterative sequence profile searches to collect HrpA and related sequences, against the non-redundant protein database (nr) from the National Center for Biotechnology Information (NCBI)^36^ clustered down to 50% identity (nr50). Clustering is based on percentage or bit-score similarity, performed using MMseqs (RRID:SCR_008184),^37^ and clustering parameters changed according to the clustering objective. The parameters of 80% coverage and 70% of identity were used to remove redundancy, and the parameters of 80% coverage and e-value of 10^−3^ were used to create homolog groups. Multiple alignments were produced with MAFFT using the local-pair algorithm combined with –maxiterate 3000 –op 1.5 –ep 0.2 (RRID:SCR_011811)^38^ or with FAMSA at default settings (RRID:SCR_021804).^39^ Domains in HrpA with known homology were annotated using a database of domain sequence profiles including pfam A models (RRID:SCR_004726).^40^

### Bacterial strains and plasmids

Deletion strains of MG1655 were constructed using one-step genomic replacement with a PCR fragment with λ Red recombinase.^41^ Gene deletions and the endogenous epitope-tagged HrpA were verified by PCR and sequencing.

The NanoLuc–BleR reporter constructs pKS-secM-short and pKS-Arg12 were expressed from plasmids containing an AmpR marker and a p15A origin of replication. To construct the Crp collision reporters, the first 39, 99, and 210 bases of the *crp* coding region were amplified from MG1655 genomic DNA. PCR was also used to add the sequence encoding the Arg12 stalling motif in frame immediately downstream of the *crp* fragment. The PCR products were inserted into EcoRI- and BglII-digested pKS-Arg12 to produce the pCRP39, pCRP99, and pCRP210 reporter plasmids using Gibson assembly. The names of the plasmids and numbers in the text represent the distance from the AUG to the stall site (the first Arg codon) in each reporter.

To construct the plasmid pAC05 encoding wild-type HrpA-FLAG, PCR was used to amplify the *hrpA-FLAG* gene, its predicted promoter, and its predicted terminator from AC016 genomic DNA. The AC016 strain expresses endogenous HrpA with a C-terminal FLAG tag. PCR was also used to amplify the origin and TetR regions from pBR322 to make the backbone. The PCR products were assembled using Gibson assembly. Site-directed mutagenesis was performed on pAC05 to produce pAC06 and pAC10 encoding the D197A and K106A HrpA mutants, respectively.

To construct the plasmid pAC08 encoding the HrpA^1–783^ HrpA mutant, the N-terminal and terminator regions of the *hrpA-FLAG* gene were amplified from AC016 and assembled with the pBR322 backbone by Gibson assembly. To construct the plasmid pAC21 encoding the HrpA^1–1179^ mutant, the *hrpA-FLAG* terminator region was amplified from pAC05 and inserted into EcoRI and BstXI-digested pAC05. The HrpA^784–1300^ mutant was expressed as a C-terminal MBP fusion from the plasmid pAC16. The MBP gene was amplified from p1M. PCR was also used to amplify all regions of pAC05 other than HrpA’s first 783 codons. The PCR products were assembled with Gibson assembly.

For overexpression and purification of HrpA, we amplified the *hrpA* gene from genomic DNA of MG1655, adding a 6xHis tag and HRV 3C cleavage site to the N-terminus of HrpA using nested PCR primers. This amplicon was inserted into pET24b digested with NdeI and XhoI using Gibson assembly.

### Growth assays

Cells were grown overnight at 37 °C in liquid LB. The overnight cultures were diluted 100-fold in fresh LB and were cultured at 37 °C to log phase (OD_600_ of approximately 0.5). Cells were diluted to prepare fivefold serial dilutions starting from OD_600_ = 0.005. Subsequently, 1.5 μL of the diluted cultures was spotted on LB plates with or without erythromycin (100 µg mL^−1^) or chloramphenicol (2.5 µg mL^−1^). Plates were then incubated at 37 °C.

### Western blots

Cells were grown in LB + ampicillin (50 mg/L) to OD_600_ = 0.5, harvested by centrifugation, resuspended in 12.5 mM Tris pH 6.8 with 4% SDS and lysed by heating to 90 °C for 10 min. Then, 5x loading dye (250 mM Tris pH 6.8, 20% glycerol, 30% β-mercaptoethanol, 10% SDS, saturated bromophenol blue) was added, and lysate was denatured at 90 °C for 10 min. Protein was separated on a 4–12% Criterion XT Bis-Tris protein gel (Bio-Rad) with XT MES buffer and was transferred to polyvinylidene difluoride membranes using the Trans-Blot Turbo Transfer system (Bio-Rad). Membranes were blocked in 5% milk for 1 h at room temperature, washed and then probed with antibodies diluted in TBS-Tween at the following dilutions: anti-Flag-HRP, 1:5,000 (Sigma); anti-RpoB, 1:1,000 (BioLegend); and anti-mouse-HRP, 1:2,000 (Thermo Fisher). Chemiluminescent signals from HRP were detected using SuperSignal West Pico PLUS Chemiluminescent Substrate (Thermo Fisher) or SuperSignal West Femto Maximum Sensitivity Substrate (Thermo Fisher) and were visualized using a BioRad ChemiDoc™ Imaging System.

### Northern blots

Cells were grown in LB + ampicillin (50 mg/L) to OD_600_ = 0.5, harvested by centrifugation and resuspended in 100 mM NaCl, 10 mM Tris pH 8.0, 1 mM EDTA pH 8.0 and 1% SDS. RNA was extracted twice with phenol (pH 4.5) at 65 °C and room temperature and subsequently by chloroform extraction. The RNA in the aqueous layer was then precipitated with isopropanol and 0.3 M sodium acetate (pH 5.5), washed with 80% ethanol and resuspended in water. The purified RNA was separated on a 1.7% agarose–formaldehyde denaturing gel and was then transferred to a nylon membrane (Hybond-N+, Cytiva) in 10× SSC buffer using a model 785 vacuum blotter (Bio-Rad). RNA was cross-linked to the membrane using an ultraviolet (UV) cross-linker (Stratagene). Pre-hybridization and hybridization were performed in PerfectHyb Plus Hybridization Buffer (Millipore Sigma). The RNA was probed with 50 nM 5ʹ-digoxigenin-labelled DNA oligonucleotides (IDT). Digoxigenin was detected with anti-digoxigenin-AP antibodies diluted 1:1,000 (Millipore Sigma). Chemiluminescent signals from alkaline phosphatase were detected with CDP-Star (Millipore Sigma) and were visualized using a BioRad ChemiDoc™ Imaging System.

### 5ʹ RACE

RNA was extracted as described above. DNA contamination was depleted by treatment with RQ1 DNase (Promega). Purified RNA (5 μg) was 5ʹ phosphorylated by incubating with T4 polynucleotide kinase (PNK, NEB) in 1 mM ATP at 37 °C for 30 min, after which PNK was denatured by heating to 75 °C for 10 min. The RNA adaptor KS_5RACE_linker was ligated to the 5ʹ end of the RNA by incubating with T4 RNA ligase 1 (NEB) in 1 mM ATP and 15% PEG 8000 at 25 °C for 3 h. The ligated samples were purified using 2.2 volumes of RNAclean XP (Beckman). First-strand cDNA was synthesized using the KS_5RACE_RT primer and SuperScript III Reverse Transcriptase (Thermo Fisher) by incubating at 54 °C for 60 min; afterwards, the reverse transcriptase was denatured by heating to 85 °C for 5 min. The denatured RT products were used directly in the first PCR. The first PCR was performed using Phusion High-Fidelity DNA Polymerase (NEB) and primers KS_5RACE_F1 and KS_5RACE_R1 with the following program: 30 s at 98 °C; 25 cycles of 10 s at 98 °C, 10 s at 65 °C and 60 s at 72 °C; and 5 min at 72 °C. Products from the first PCR were purified using DNA Clean & Concentrator-5 columns (Zymo Research). The second PCR was performed using Phusion High-Fidelity DNA Polymerase (NEB) and primer NI-NI-2 in addition to one of our custom index primers for 5ʹ RACE (Supplementary Table 1) with the following program: 30 s at 98 °C; 25 cycles of 10 s at 98 °C, 10 s at 65 °C and 60 s at 72 °C; and 5 min at 72 °C. Products from the second PCR were purified using DNA Clean & Concentrator-5 columns (Zymo Research), analyzed using a BioAnalyzer high-sensitivity DNA kit (Agilent) and sequenced on a MiSeq Nano instrument (Illumina).

The 5ʹ-RACE data were analyzed using custom scripts written in Python 2.7. The RNA adaptor sequence TGCCCGAGTG was removed from the 5ʹ end of the reads using Cutadapt (https://doi.org/10.14806/ej.17.1.200). Reads without the RNA adaptor sequence were discarded. The reverse primer sequence GCGGTCGAGTTCTGGACCGA from the second PCR was removed from the 3ʹ ends of the reads using cutadapt. The processed reads were aligned to the short SecM reporter plasmid sequence using bowtie.^42^ The 5ʹ ends of the mapped reads were counted and normalized as RPM mapped reads for the sequencing depth of each library.

### Ribosome profiling

Ribosome profiling was performed as described.^21^ In brief, cells expressing the pKS-secM-short reporter were grown in 100 mL MOPS defined media (Teknova) to OD 0.5 and harvested by spraying directly into liquid nitrogen. Cells were lysed using a Spex 6870 freezer mill, and the lysates were centrifuged to pellet cell debris. Supernatants were then layered over sucrose cushions and ultracentrifuged to pellet the ribosomes. Pellets were resuspended, treated with MNase to digest unprotected mRNA, and centrifuged through sucrose gradients. Fractions containing monosomes were then collected and the RNA was extracted. Universal miRNA cloning linker (NEB) was ligated to RNAs 15-45 nt in length, the RNAs were reverse transcribed, and resulting cDNAs 15-45 nt in length were circularized. rRNA was depleted prior to PCR and sequencing. Resulting sequence reads were filtered, trimmed, aligned, and used to calculate ribosome density on reporter mRNA.

### Polysome profiling

Cells were cultured at 37 °C in 500 mL of LB and antibiotics where appropriate to OD_600_ = 0.5. For samples with antibiotic treatment, cells were cultured to OD_600_ = 0.45 and subsequently treated for 5 min with antibiotics at the concentrations indicated in the figures. Cells were then harvested by filtration using a Kontes 99-mm filtration apparatus with a 0.45-μm nitrocellulose filter (Whatman) and were flash frozen in liquid nitrogen. Cells were then lysed in lysis buffer (20 mM Tris pH 8.0, 10 mM MgCl_2_, 100 mM NH_4_Cl, 5 mM CaCl_2_, 100 U/mL DNase I, 1 mM chloramphenicol) using a Spex 6870 freezer mill with five cycles of 1 min grinding at 5 Hz and 1 min cooling. Lysates were centrifuged at 20,000*g* for 30 min at 4 °C to pellet the cell debris. Sucrose gradients of 10–50% sucrose were prepared using a Gradient Master 108 (Biocomp) with gradient buffer (20 mM Tris pH 8.0, 6 mM MgCl_2_, 100 mM NH_4_Cl, 2 mM DTT). Then, 5–40 AU of *E. coli* lysate was loaded on top of the sucrose gradient and was centrifuged in an SW 41 rotor at 35,000 rpm. for 2.5 h at 4 °C. Fractionation was performed on a Piston Gradient Fractionator (Biocomp). To process each fraction for western blots, proteins were precipitated in 10% TCA. Afterwards, the pellets were washed twice in ice-cold acetone, vacuum dried briefly, resuspended in 5x loading dye and neutralized with Tris-HCl pH 7.5.

### Purification of HrpA and inactive HrpA mutants (K106A, D197A, HrpA^1–783^, HrpA^1–1179^)

pET24b plasmids encoding HrpA or the HrpA mutants with an N-terminal 6xHis-tag and an HRV 3C cleavage site were introduced into *E. coli* strain BL21 (DE3). Cells were grown in 1 L of LB to mid-log phase (OD_600_ = 0.5) at 37 °C and were induced with 0.125 mM IPTG at 18 °C for 20 h. Cells were harvested by centrifugation at 4,000*g* at 4 °C for 5 min, resuspended with buffer A (50 mM Tris-HCl pH 8, 500 mM NaCl, 2 mM β-mercaptoethanol, 0.02% NP-40, 0.2 mg mL^−1^ lysozyme, 2 µL ml^−1^ RQ1 DNase, 1:1,000 dilution of protease inhibitor (pill per mL), 25 mM imidazole) and lysed by sonication (Branson 250 Sonicator). Cell debris was removed by centrifugation at 30,000*g* at 4 °C for 30 min. The cleared lysate was then incubated with 2 mL of prewashed PureCube Ni-NTA agarose (Cube Biotech) for 1 h. Afterwards, the beads were washed with 20 column volumes (CVs) buffer B (50 mM Tris-HCl pH 8, 500 mM NaCl, 25 mM imidazole) and twice with 10 CVs buffer C (10 mM Tris-HCl pH 8, 200 mM NaCl, 25 mM imidazole). Samples were then eluted with 5 CV buffer D (10 mM Tris-HCl pH 8, 200 mM NaCl, 250 mM imidazole, 1 mM DTT).

Next, the eluate was either diluted to 100 mM NaCl with Tris-HCl pH 8 (wild-type, K106A, and D197A) or kept undiluted (HrpA^1–783^ and HrpA^1–1179^). The samples were loaded onto a HiTrap™ Heparin HP column (Cytiva) equilibrated with 10 mM Tris-HCl pH 8, 100 mM NaCl, and 1 mM DTT. The protein was eluted with a linear gradient from 100 to 1,500 mM NaCl. Fractions containing protein were concentrated to 1 mL using an Amicon 100,000 MWCO (or 30,000 MWCO for HrpA^1–783^ and HrpA^1–1179^) device and further purified by size exclusion chromatography on a 10/30 Superdex™ 200 column (GE Healthcare) in 10 mM Tris-HCl pH 8, 200 mM NaCl, and 1 mM DTT. The HrpA-containing fractions were again concentrated and stored at −80 °C.

### Polysome enrichment for in vitro splitting

Δ*hrpA* cells were cultured at 37 °C in 500 mL of LB to OD_600_ = 0.5. For samples with antibiotic treatment, cells were cultured to OD_600_ = 0.45 and subsequently treated for 5 min with 5 or 500 µg mL^−1^ chloramphenicol. Cells were then harvested by filtration using a Kontes 99-mm filtration apparatus with a 0.45-μm nitrocellulose filter (Whatman) and were flash frozen in liquid nitrogen. Cells were then lysed in lysis buffer (20 mM Tris-HCl pH 8.0, 150 mM MgCl_2_, 100 mM NH_4_Cl, 5 mM CaCl_2_, 100 U mL^−1^ DNase I, 0.4% Triton X-100, 0.1% NP-40) using a Spex 6870 freezer mill with five cycles of 1 min grinding at 5 Hz and 1 min cooling. Lysates were centrifuged at 20,000*g* for 30 min at 4 °C to pellet the cell debris. Supernatants were then layered over 2 mL sucrose cushion (1.45 M sucrose, 20 mM Tris-HCl pH 8, 500 mM NH_4_Cl, 10 mM MgCl_2_, 0.5 mM EDTA pH 8) and centrifuged in a TLA 100.3 rotor at 83,000 rpm for 20 min at 4 °C. Pellets were resuspended in 20 mM Tris-HCl pH 8, 6 mM MgCl_2_, 100 mM NH_4_Cl, and 5 mM CaCl_2_ and stored at −80 °C.

### In vitro splitting of enriched polysomes

25 pmol of enriched polysomes were incubated with 1 mM ATP, 250 pmol purified IF3, purified HrpA, and 4 µg mL^−1^ purified creatine kinase with 12 mM phosphocreatine (for regenerating ATP) for 1 hr at 25 °C. Sucrose gradients of 10–50% sucrose were prepared using a Gradient Master 108 (Biocomp) with gradient buffer (20 mM Tris pH 8.0, 6 mM MgCl_2_, 100 mM NH_4_Cl, 2 mM DTT). Then, the splitting reactions were loaded on top of the sucrose gradient and centrifuged in an SW 41 rotor at 35,000 rpm for 2.5 h at 4 °C. Fractionation was performed on a Piston Gradient Fractionator (Biocomp).

### PURExpress in vitro translation and isolation of polysomes

To obtain a template for *in vitro* translation, PCR was used to amplify linear DNA from pKS-secM. Resulting templates encoded a T7 promoter, ribosome binding site, the NanoLuc-SecM coding region, and 370 nt downstream of the SecM stalling motif. Ribosome/nascent chain complexes (RNCs) were generated with a PURExpress In Vitro Protein Synthesis Kit (New England Biolabs, E6800S, transcription and translation coupled) using 400 ng of template DNA for each 25 μL reaction. Five reactions were incubated at 37 °C for 1 h and were subsequently combined and loaded on sucrose gradients (20 mM Tris pH 8.0, 6 mM MgCl_2_, 100 mM NH_4_Cl, 2 mM DTT; 10–50% sucrose) and spun in an SW 41 Ti rotor (Beckman Coulter) at 35,000 r.p.m for 2.5 h at 4 °C. The gradient was fractionated at a BioComp Gradient Station IP using a Triax flow cell for UV measurement. The polysome peak fractions were collected and pelleted by centrifugation in a TLA 100.3 rotor (Beckman Coulter) at 95,000 r.p.m for 2 h at 4 °C. After resuspension in RNC buffer (20 mM Tris-HCl pH 8, 10 mM MgCl_2_, 100 mM NH_4_Cl, and 5 mM CaCl_2_), samples were frozen in liquid nitrogen and stored at −80 °C.

### In vitro splitting of PURExpress polysomes

20 pmol of polysomes were reacted with 20 pmol purified HrpA, 1 mM ATP, 200 pmol purified IF3, 4 µg mL^−1^ creatine kinase and 12 mM phosphocreatine for 1 h at 25 °C.

### Purification of HrpA for cryo-EM

HrpA was cloned into a pBAD vector following a N-terminal His-tag and 3C cleavage site and overexpressed in BL21(DE3). 3 x 2 L LB with ampicillin were inoculated and grown at 125 rpm, 37 °C to OD600 = 1.2, then induced with 0.2 % arabinose overnight at 16 °C. Cells were harvested by centrifugation in a F9-6 x 1000 LEX rotor at 4500 rpm for 5 min, washed in cold 1X PBS, and resuspended in cold lysis buffer (50 mM HEPES-NaOH pH 8.0, 500 mM NaCl, 100 mM PMSF, 2 mM BME, 1/50 pill/ml cOmplete™, EDTA-free Protease Inhibitor Cocktail (Roche)). DNaseI was added to resuspended cells to a final concentration of 10 μg/mL prior to lysis by one pass through a cell disruptor (Constant Systems) at 18 kPsi at 4 °C. Lysate was supplemented with 0.02% (v/v) Igepal and clarified by centrifugation in a SS-34 rotor at 18,000 rpm for 20 min at 4°C. Half of this lysate was frozen in liquid N2 and stored at −80 °C until use for the purification. Lysate was thawed, supplemented with 20 mM Imidazole and incubated for 30 min, rotating at 4°C, with 2 mL Ni-NTA beads (4 mL slurry), which had been pre-equilibrated with 50 mM HEPES-NaOH pH 8.0, 500 mM NaCl, 5% glycerol, 2 mM BME, 20 mM Imidazole, 0.02% (v/v) Igepal. Beads were then washed by gravity flow with 20 mL Wash buffer I (50 mM HEPES-NaOH pH 8.0, 500 mM NaCl, 2 mM BME, 50 mM Imidazole, 0.02% (v/v) Igepal, 1/1000 pill/ml protease inhibitor cocktail), 2 x 20 mL Wash buffer II (as wash buffer I without protease inhibitor) followed by one wash with 10 mL Wash buffer III (50 mM HEPES-NaOH pH 8.0, 500 mM NaCl, 5% glycerol, 0.02% Igepal, 2 mM β-mercaptoethanol). Elution was performed by incubation with 3 mL Wash Buffer III + 500 mM Imidazole for 10 min. HrpA elution fractions were concentrated to 1 ml in an Amicon ultracel concentrator with a MW cuttoff of 100 kDa and subjected to size exclusion chromatography on a Superdex 200 Increase column with S200 buffer (50 mM HEPES-NaOH pH 8.0, 500 mM NaCl, 5% glycerol, 1 mM DTT).

### E. coli in vitro translation and isolation of collided disomes

VemP stalled RNCs were prepared essentially as previously described.^9^ mRNA encoding VemP peptide without N-terminal signal sequence, preceded by an N-terminal cleavable His-tag and a Flag-tag, was prepared by PCR amplification, DNA purification, *in vitro* transcription and precipitation by LiCl. Plasmid pHK060 was used with primers 5’-cgatgcgtccggcgtagaggatcg-3’ and 5’GGTTATAATGAATTTTGCTTATTAACcctttcgggctttgttagca-3’ to generate a template for *in vitro* transcription, producing a 698 nt long mRNA, of which 147 nt are downstream of the A-site codon in the stalled ribosome. Ribosome nascent chain complexes were generated using the PURExpress In Vitro Protein Synthesis Kit (New England Biolabs, E6800S). A 100 µL reaction was used, with 90 µg mRNA and 32 U superaseIn RNase inhibitor. Reactions were incubated at 30 °C for 35 min, then loaded on a 10-50% sucrose gradient (20 mM Hepes pH 7.5, 10 mM Mg(OAc)_2_, 150 mM KOAc, 1 mM DTT) and centrifuged for 13 h in a SW40 Ti rotor (Beckman Coulter) at 19,200 rpm. Gradients were fractionated using a Biocomp Gradient Station ip and the disome peak was collected. Disomes were pelleted in a TLA110 rotor (Beckman Coulter) for 45 min at 100,000 rpm, 4 °C, and resuspended in RNC buffer (25 mM Hepes/KOH pH 7.5, 150 mM KOAc, 10 mM Mg(OAc)_2_, 1 mM DTT).

### Sample preparation for cryo-EM

A 20 µL reaction with 2.8 pmol ribosomes (0.14 A_260_ units), 11-fold excess (1.5 µM) HrpA, and 1 mM AMP-PNP in RNC buffer was incubated for 5 min at 30 °C. Nikkol was added to a final concentration of 0.05 % (w/v), and 3.5 µL sample applied to 2 nm carbon coated Quantifoil R3/3 holey grids, which were vitrified in liquid ethane using a Vitrobot mark IV (FEI) with a blotting time of 3 s and wait time of 45 s.

### Cryo-EM data acquisition and data processing

Data were collected with EPU software (v3.7) a Titan Krios electron microscope at 300 kV with a SelectrisX energy filter using a Falcon 4i detector, with a defocus range 0.5 µm - 3.5 µm, total dose of 40 e^−/Å2^ and nominal pixel size of 0.727 Å. All 40 frames were gain corrected, aligned and summed using MotionCor2,^43^ and CTF parameters were estimated using CTFFind4.^44^ After visual inspection 37,100 micrographs were subjected to Laplacian-of-Gaussian based auto-picking in RELION 5.0 to pick 70S ribosomes. 2,008,477 particles were extracted with a binning factor of 6 and box size of 84 pixels and subjected to 2D classification in CryoSPARC (v4.4.0). 1,753,301 ribosomal particles were re-extracted with a binning factor of 4 and box size of 140 and refined against a 70S ribosome in RELION. Initial 3D classification was performed with a mask covering the positions of neighboring ribosomes (either stalled or collided). The class in which the stalled ribosome, occupied with A/A and P/P-tRNAs, was centered (462,831 particles) was refined, extracted with a larger box size of 160 pixels and subjected to a further round of refinement. Focused 3D classification with a mask around HrpA resulted in a class with an elongated density spanning over the 30S head of the stalled ribosome and connecting over to the 30S of the collided one (Fig. 3A; Fig. S6). This was further sub-classified with a mask around the connector and hook domains giving rise to one class containing one HrpA molecule (Class Ia, 41,265 particles) and a second class with additional density for a hook/α+β domain of a second HrpA molecule (Class IIa, 112,106 particles). Class Ia was refined, extracted unbinned with a box size of 600 pixels and pixel size of 0.727 Å. CTF refinement was carried out, estimating parameters for beam tilt and trefoil, anisotropic magnification, and astigmatism (per micrograph) and defocus (per particle), which was followed by 3D refinement. Following this, the whole disome was extracted with a box of 240 pixels and 2.181 Å/pixel, refined, and further classified with a mask around the L1 stalk. Refinement was then performed with unbinned particles with a box of 700 pixels. This yielded HrpA-bound disome classes with L1 of the collided disome either in the “out” or “in” position, and with the collided ribosome occupied either by canonical A/A and P/P tRNAs or by hybrid A/P and P/E tRNAs respectively. These two classes (Class I, L1-out; 17,182 particles, and L1-in; 20,445 particles) were then refined in CryoSPARC to a resolution of 3.1 Å and 3.2 Å respectively. A local refinement was performed on Class I using a mask around the helicase domain of HrpA. Note that whenever particles were re-extracted with a larger box size, particles at the micrograph edges were discarded.

Class IIa, which contains extra density for a second HrpA molecule, was subclassified with a mask around the entire helicase and β-meander domains of HrpA. This yielded one class with extra tube-like density reaching alongside the HB2 and β-meander domains of HrpA1, likely accounted for by further domains of the second HrpA molecule, and a further five classes with the helicase in slightly different positions.

### Model building

The structure of the VemP-stalled collided disome (PDB-IDs 7qg8 and 7qgh)^9^ served as the initial model for the HrpA-bound disome. HrpA was modelled based on an AlphaFold2 prediction^24^ and a crystal structure of the N-terminal helicase domain complexed with RNA (PDB-ID 6zww).^15^ The crystal structure was split into two parts, one containing the two RecA domains along with the RNA, and the other containing the HB2 domain. Both parts were individually docked into the locally refined map for the HrpA helicase (see Fig. S6) and ISOLDE was used to fix clashes.^45^ WH, OB and β-meander domains were positioned based on the fitted HB2 domain and matched the density well without further adjustments. The remaining HrpA structure (from Asp741 to 1300) was fitted based on the Alphafold2 model, which was split into regions covering the HB2, α+β and Hook domains. These domains were then individually docked into the map and adjusted with Coot^46^ and ISOLDE^45^ in ChimeraX (v1.71).^47^

Density for mRNA was visible throughout the VemP-stalled collided disome and emerging into the HrpA helicase core. Based on mRNA from 7qgh (14 nts modelled), we were able to model 57 nucleotides spanning from the E-site of the collided ribosome via the stalled ribosome into the HrpA helicase. Where density became unclear between mRNA entry site and helicase, the sequence was modelled as poly-U. Since the frame of mRNA could be identified in the collided ribosome, A-site and P-site tRNAs were altered to match the sequences of Met and Ala tRNA respectively.

In addition to 7qg8 and 7qgh, models were improved for bL31, where a helix covering residues 55-66 (visible in PDB 7k00)^48^ was docked and for bS18 in the collided ribosome (based on 7k00). Moreover, clear density was visible for the stalk base, where nucleotides 1045-1106 of 23S rRNA, uL10 and uL11 of the stalled ribosome were modelled based on 8pkl.^23^ Ribosomal protein bS1 was omitted in the final model. The model for the second HrpA copy (HrpA2) was generated by rigid-body docking the Hook domain as well as part of the α+β domain (residues 1011-1300) into the respective density of ClassII (see Fig. S4). Following rigid-body fitting and real space refinement in Coot, the complete model (except for the HrpA helicase module) was then refined using Phenix (v.1.12-4487).^49^

### Post-processing and map display

To generate the composite map for the HrpA-bound disome as displayed in Figure 3, the unmodified cryo-EM map (i.e., before post-processing) for Class I (disome bound to one Hrp1 copy; see Fig. S4) was segmented into isolated densities for the disome and HrpA Hook/α+β and β-meander/HB2 domains. HrpA domains were then gaussian low-pass filtered in ChimeraX (standard deviation of 0.8 for Hook/α+β and 1.0 for β-meander/HB2). Density for the HrpA helicase domain was isolated from the locally refined map (Fig. S6). To generate the composite map for the disome bound to two HrpA copies as displayed in Fig. S10, the refined map was low-pass filtered according to local resolution. The map was then segmented into isolated densities for the disome, the first HrpA, the hook domain of the second HrpA and remaining extra density. All figures containing models and densities were prepared using UCSF ChimeraX (version 1.71).^47^

