## Supplemental Figures for "The RNA helicase HrpA rescues collided ribosomes in *E. coli*"

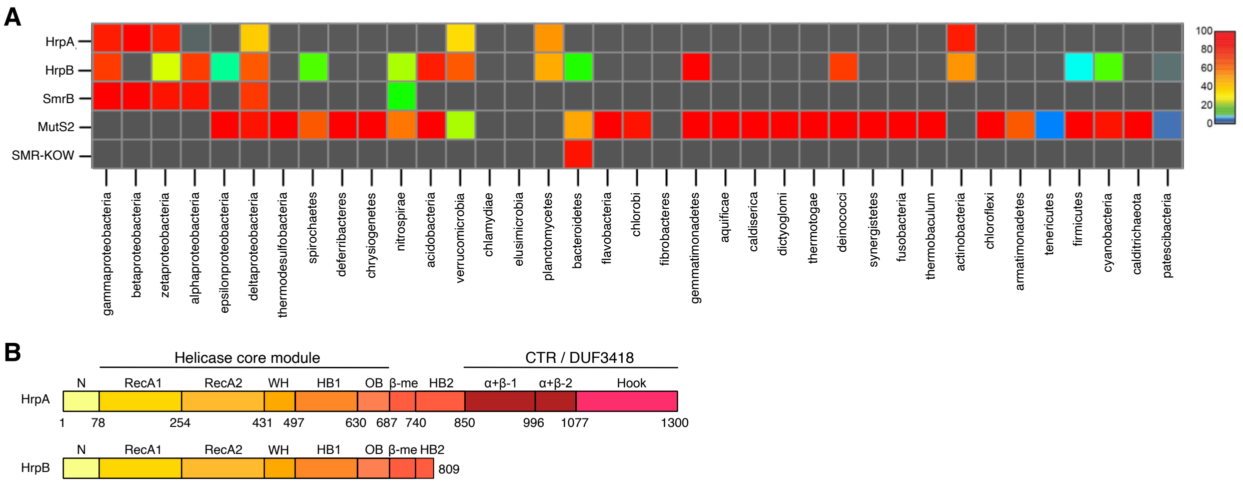

**Fig. S1. HrpA is a conserved DExH-box protein.** (**A**) Heatmap showing the percentage of genomes in each bacterial phylum encoding HrpA, HrpB, and SMR domain-containing proteins. (**B**) Domain architectures of *E. coli* proteins HrpA and HrpB, which lacks the C-terminal region found in HrpA.

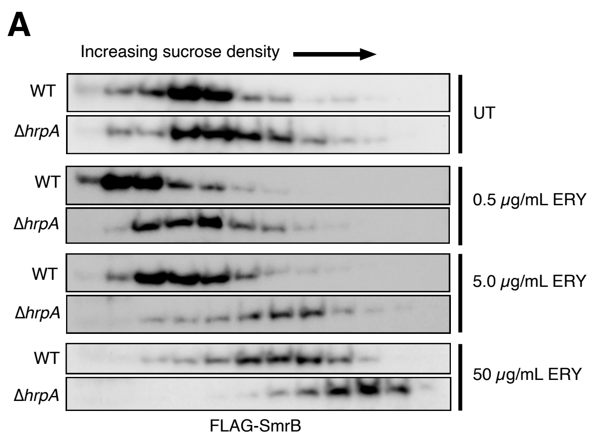

**Fig. S2. The effects of HrpA on SmrB migration in sucrose gradients**. Distribution of FLAG-SmrB in sucrose gradients observed with anti-FLAG antibodies.

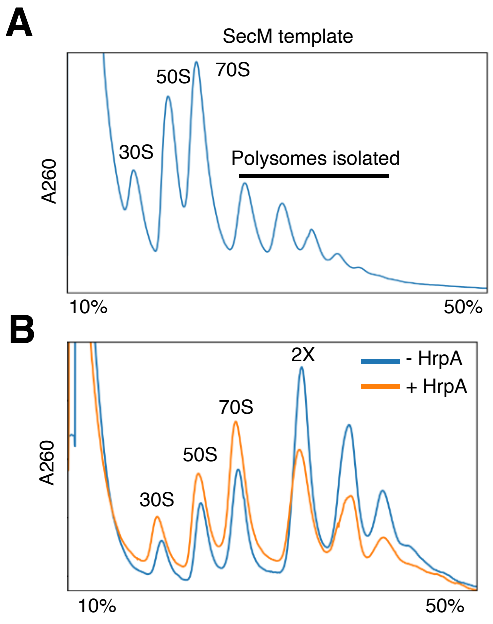

**Fig. S3. HrpA splits ribosomes stalled on mRNA in vitro.** (**A**) Polysome traces from PURE reactions with the SecM stalling reporter. (**B**) Polysome traces from *in vitro* reactions with purified polysomes from SecM reporters with 300 nt downstream of the stall site.

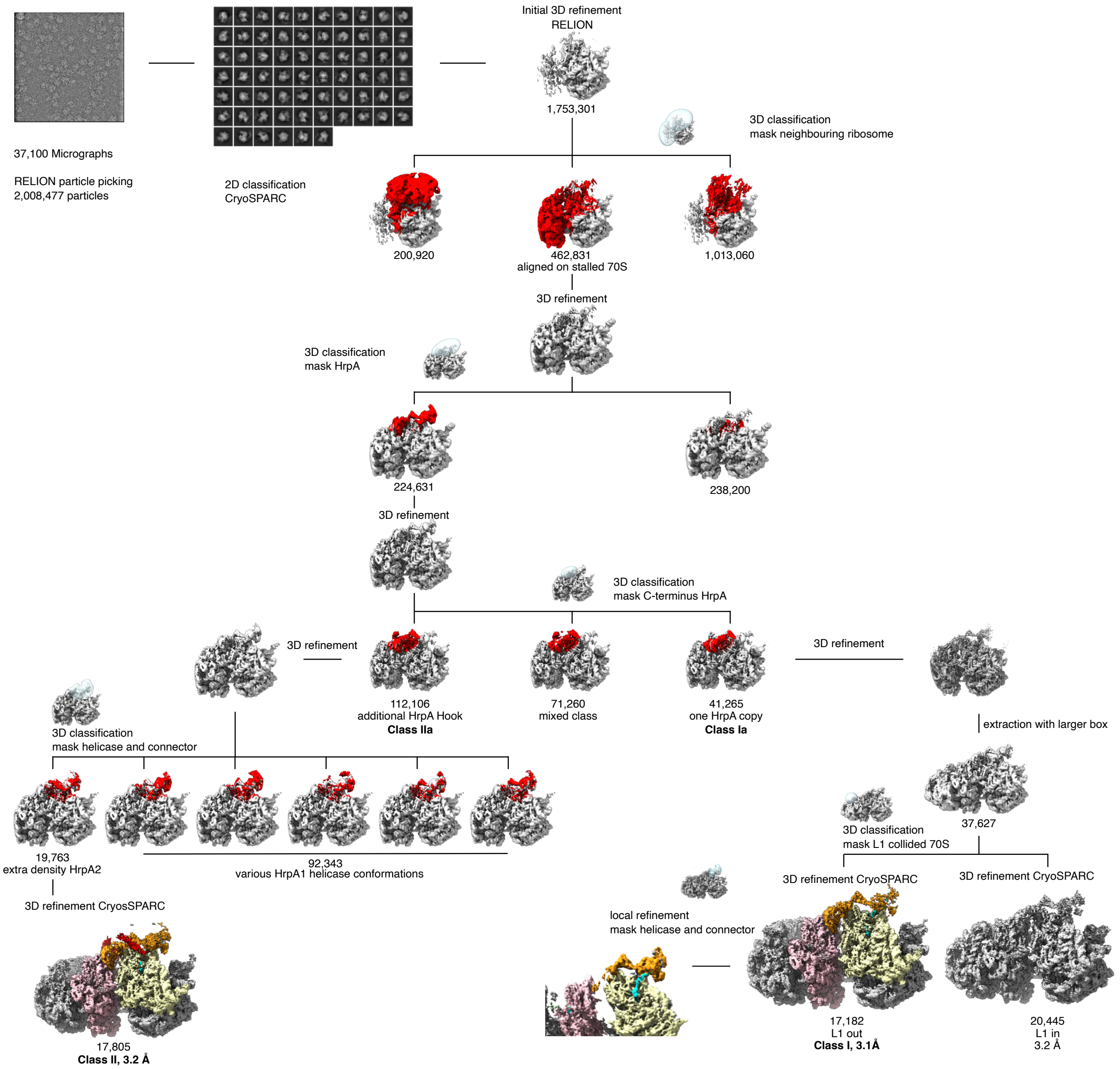

**Fig. S4. Cryo-EM data analysis and classification of HrpA-disome complexes.** From a total of 37,100 micrographs, 2,008,477 particles were picked using RELION AutoPick and used for 2D classification, which yielded a total of 1,753,301 70S ribosomal particles. After initial refinement of a 70S ribosome, 3D classification was performed with a mask covering the positions of a neighbouring ribosome (either the stalled or the collided 70S) in the disome. This yielded two stable disome classes, with one aligned on the stalled 70S and the other aligned on the collided 70S. The larger class (462,831 particles) showed extra-ribosomal density for HrpA and was further sorted by focused classifications: first using a mask for HrpA, yielding enriched HrpA extra density, and then using a smaller mask covering only the Hook and connector domains. This resulted in three classes, one (Class Ia) showing one copy of HrpA bound to the disome (41,265 particles), one showing an additional second Hook density for HrpA (112,106 particles, Class IIa) and one mixed class (71,260 particles) that was not further classified.

Class Ia was further split by focused classification using a mask on L1 of the collided ribosome. This resulted in an HrpA-bound disome with L1 of the collided ribosome in the “out”-position together with A/A and

P/P site tRNAs (Class-I; 17,182 particles) and with L1 of the collided ribosome in the “in” position together with hybrid state A/P, P/E tRNAs (L1 in; 20,445 particles). Subsequently, a focused refinement was performed on the HrpA helicase region of Class-I, yielding the best map of mRNA-bound HrpA (see Extended Data Fig. 7).

Class IIa was further sub-classified with a mask focusing on the HrpA helicase and connector domain, resulting in one class with clear, tube-like extra density, likely accounting for remaining domains of the second HrpA copy (Class II, 17,805 particles) and several mixed classes with various conformations of the helicase. Class I and Class II were refined to the best possible resolution, yielding final maps (3.1 Å for Class I, and 3.2 Å for Class-II).

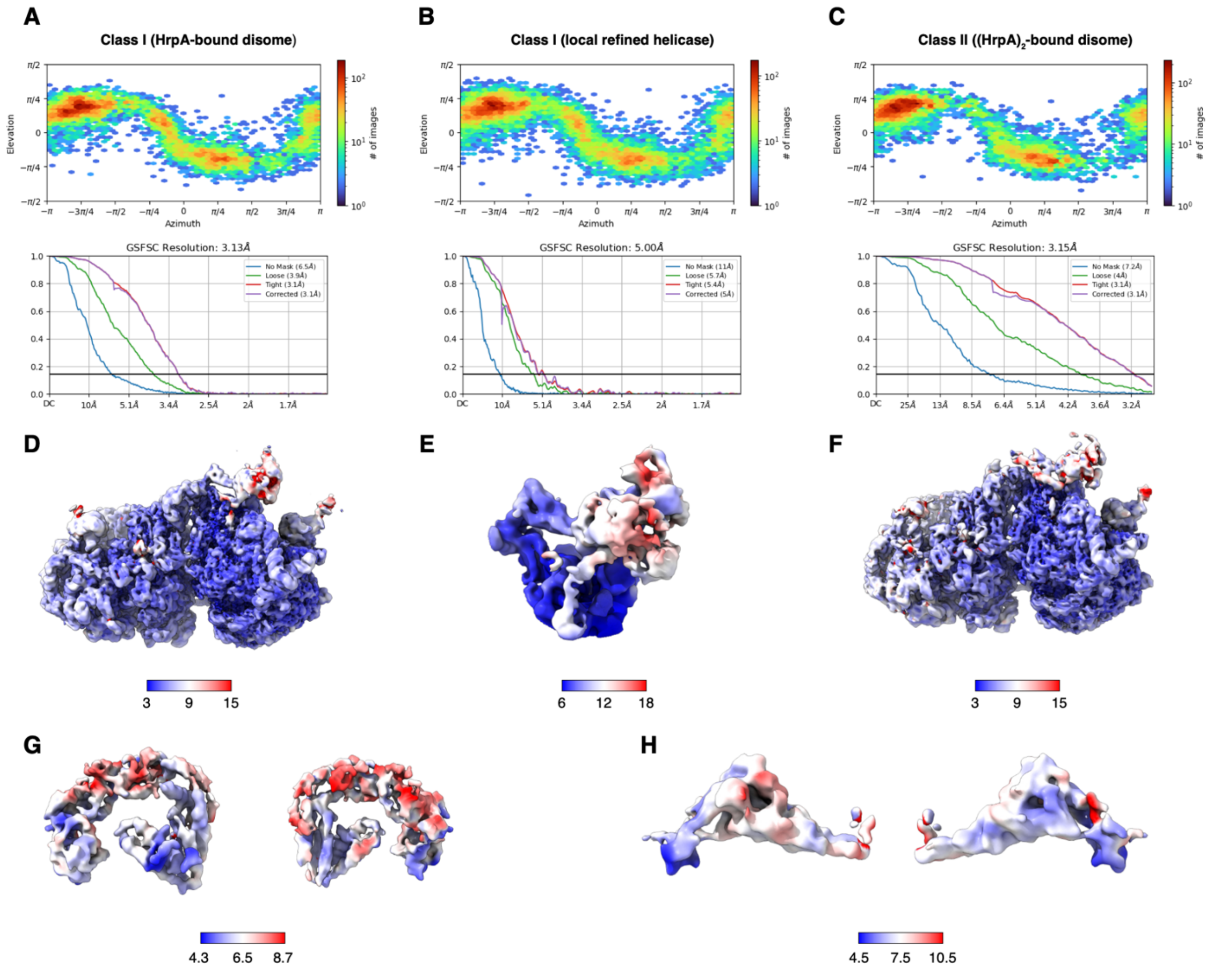

**Fig. S5. Angular distribution and resolution of HrpA-disome complexes.** (**A-C**) Angular distribution plots and Gold-standard Fourier Shell Correlation (GS-FSC) curves (obtained from CryoSPARC) for the reconstructions of Class I (HrpA-bound disome, **A**), Class I locally refined on the HrpA helicase domain (Class I-loc, **B**) and Class II ((HrpA)$_2$-bound disome, **C**) (see also fig. S4). (**D-F**) Cryo-EM maps of Class I (**D**) and Class II (**F**) were low-pass filtered and both classes as well as Class I-loc (**E**) were colored according to local resolution. (**G, H**) Isolated densities for the HrpA Hook/α+β (**G**) and β-meander/HB2 domains (**H**), gaussian low-pass filtered at a standard deviation of 0.8 (for Hook/α+β) and 1.0 (for β-meander/HB2) (as in Fig. 3A) and colored according to local resolution.

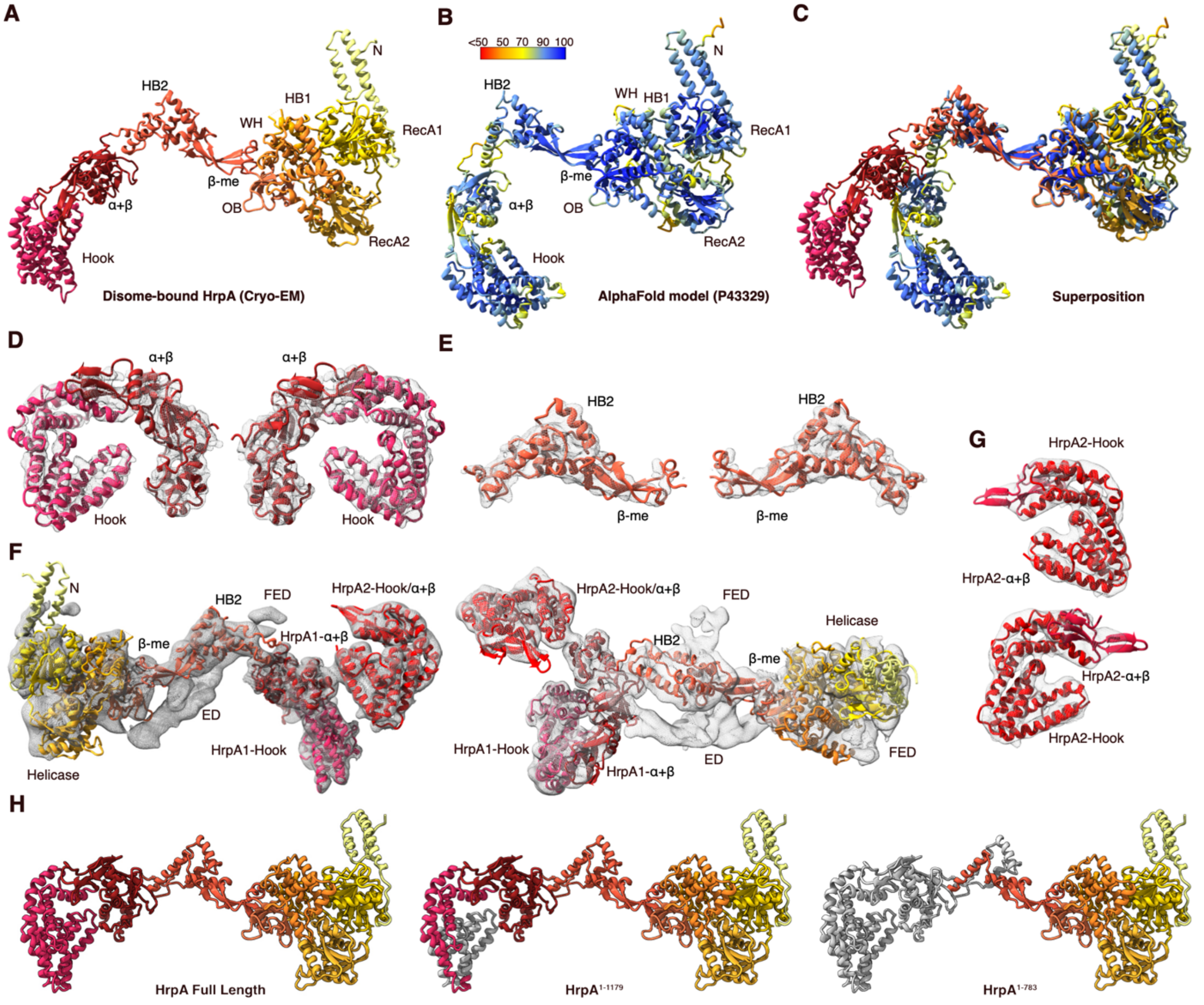

**Fig. S6. Model fitting and map quality of HrpA-disome reconstructions.** (**A**) Final model of the disome-bound HrpA complex derived from fitting the AlphaFold2 model into the cryo-EM map (Class I; see fig. S4). (**B**) AlphaFold2 model of HrpA (UniProt-ID P43389). The model is colored according to a per-model confidence score (pLDDT; from 0 to 100). Blue regions display a very high confidence (pLDDT > 90), red regions low confidence (pLDDT < 50). (**C**) Overlay of cryo-EM model with AlphaFold2 model. (**D, E**) Molecular model of HrpA Hook/α+β (**D**) and β-meander/HB2 domains (**E**) fitted into isolated densities. Densities were derived from Class I and gaussian low-pass filtered at a standard deviation of 0.8 (for Hook/α+β) and 1.0 (for β-meander/HB2) (as in Fig. 3A). Views are the same as in fig. S5G and H. (**F**) Two views showing isolated non-ribosomal density of Class II with a fitted model for disome-bound HrpA and the Hook/α+β domain of a second HrpA copy. Unassigned extra density (ED) likely accounts for the rest of the second HrpA and fuzzy extra density (FED) remains unassigned. (**G**) Two views showing the isolated density with fitted model for the Hook/α+β domain of the second HrpA copy. (**H**) Molecular model of HrpA: truncated regions in HrpA mutants used in Fig. 4 (HrpA$^{1-1179}$ and HrpA$^{1-783}$) are depicted in grey.

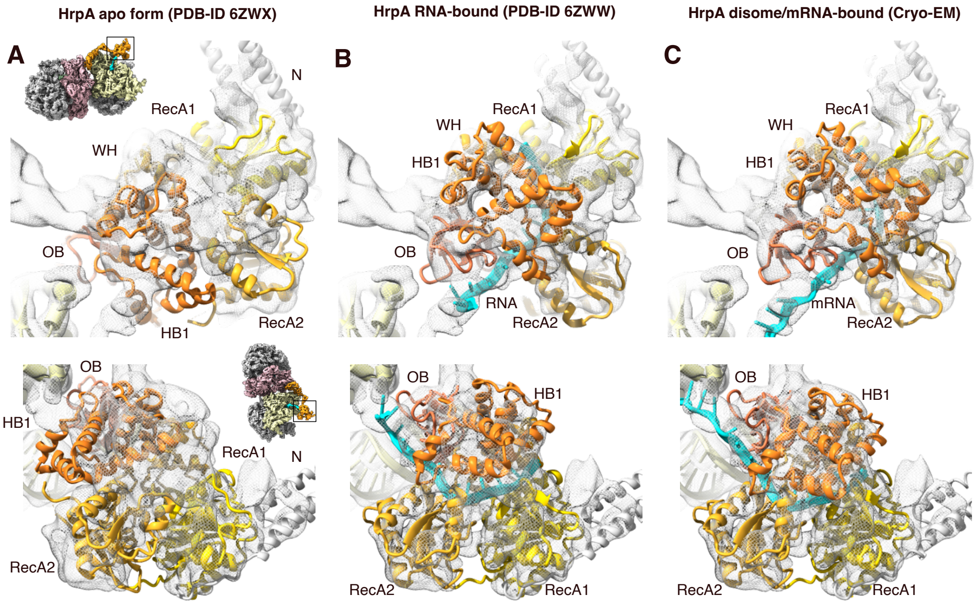

**Fig. S7. Density fits of crystal structures and disome-bound model of HrpA.** (**A-B**) Fitting of the HrpA crystal structure without (PDB-ID 6zwx, **A**) and with RNA bound (PDB-ID 6zww, **B**) into the focused refined map for the HrpA-bound disome (Class I; see fig. S4). Top panels show a view focusing on the HrpA helicase at mRNA entry channel of the stalled 70S, bottom panels focus on the HrpA helicase RecA1 and RecA2 domains. Thumbnails indicate the views and black boxes show the zoomed region. (**C**) Same as **A** and **B** with the model for the disome-bound HrpA. Maps are shown as transparent mesh.

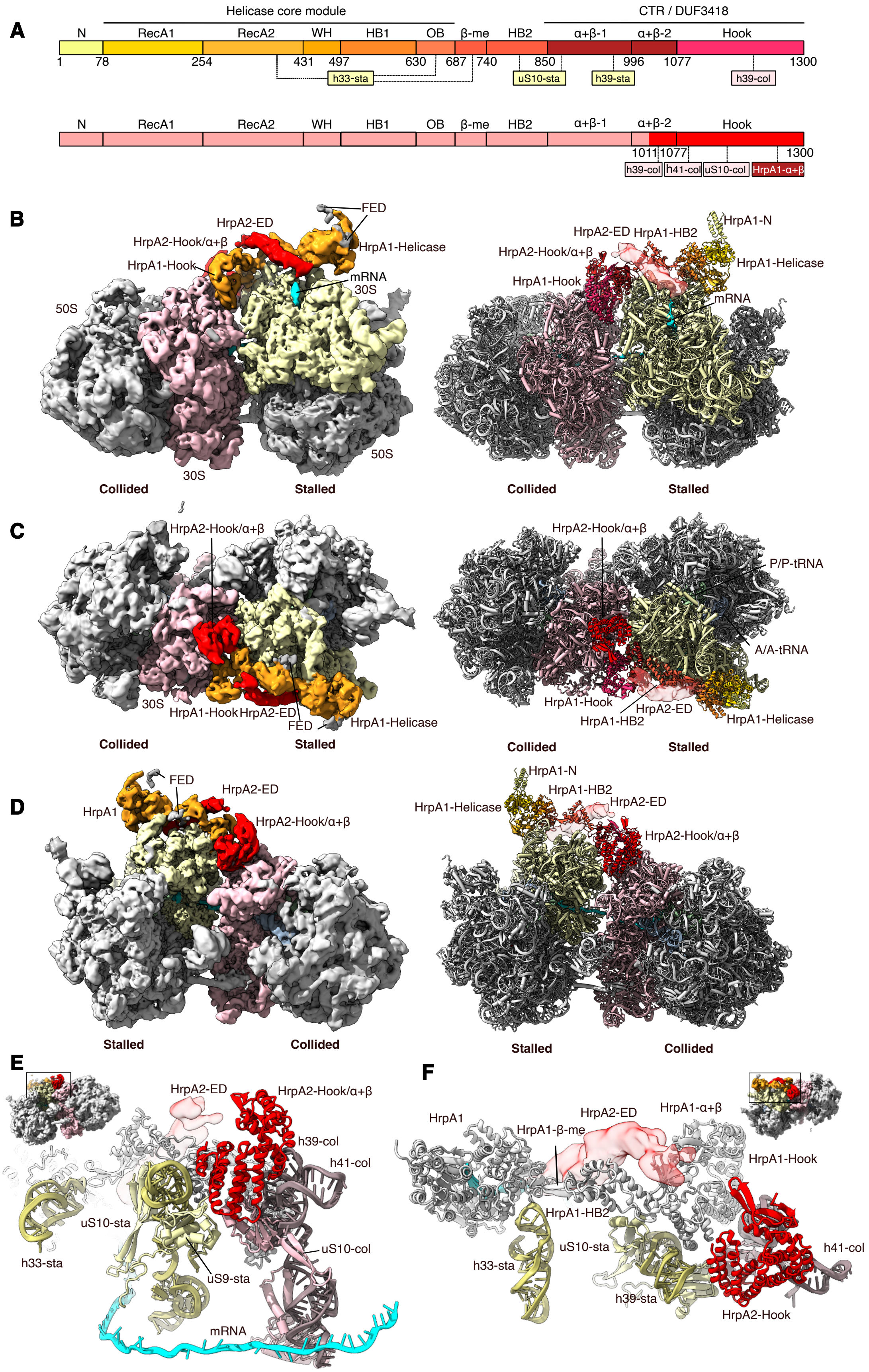

A
Helicase core module
CTR / DUF3418
N
RecA1
RecA2
WH
HB1
OB
β-me
HB2
α+β-1
α+β-2
Hook
1
78
254
431
497
630
687
740
850
996
1077
1300
h33-sta
uS10-sta
h39-sta
h39-col
1011
h41-col
uS10-col
HrpA1-α+β
B
FED
HrpA2-ED
HrpA2-Hook/α+β
HrpA1-Helicase
HrpA1-Hook
mRNA
30S
50S
Collided
Stalled
HrpA1-HB2
HrpA1-N
C
P/P-tRNA
A/A-tRNA
D
HrpA1
E
h39-col
h41-col
uS10-sta
h33-sta
uS9-sta
mRNA
F
HrpA1-β-me
HrpA1-HB2
h39-sta
HrpA2-Hook

**Fig. S8. Structure of the disome bound to two HrpA copies**. (**A**) Schematic representation as in Fig. 3D of the domain organization for the two HrpA copies (HrpA1 and HrpA2). Domains that are not visible or cannot be assigned in HrpA2 are shown in transparent red. (**B-D**) Cryo-EM density map and molecular model of the $(HrpA)_2$-disome shown as side (**B**), top (**C**) and back (**D**) views. The map was low-pass filtered according to local resolution (see fig. S4, S5 and Methods). Densities for HrpA1, HrpA2 Hook (Hook2) and HrpA2 extra density (ED) were segmented and are shown at lower contour levels as compared to the disome for clarity. The color code for HrpA domains is given in **A** and remaining fuzzy extra density (gray; FED) is indicated. (**E-F**) Molecular model the $(HrpA)_2$-disome, shown as a back view highlighting the position of Hook2 (**E**) or displaying the ED for HrpA2 adjacent to HB2, β-meander, and OB-fold domains of HrpA1 (**F**). Views are indicated in the thumbnails. Herein, the black box shows the zoomed region. HrpA1 is shown in grey and HrpA2 in red.

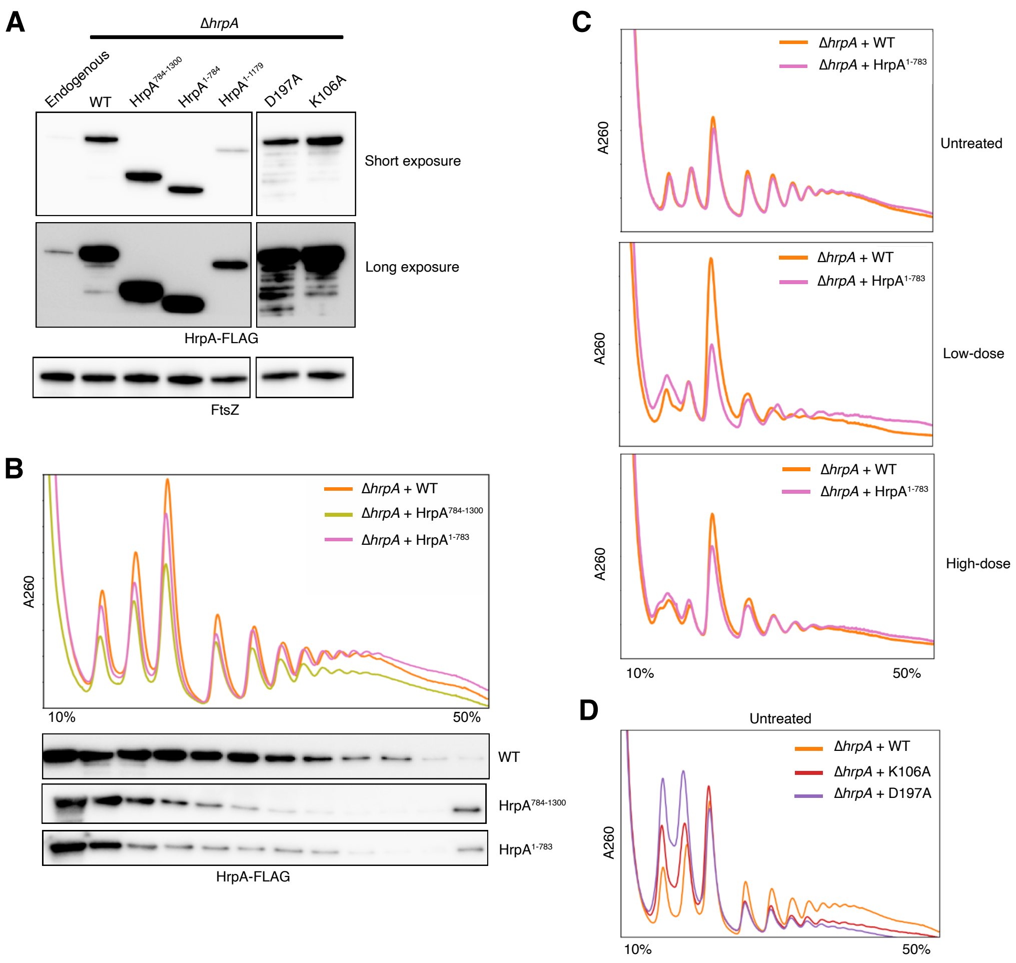

**Fig. S9. HrpA mutant protein levels and polysome traces.** (**A**) Immunoblot showing expression levels of HrpA constructs. Lane 1: Strain endogenously expressing wild-type HrpA-FLAG from its native locus. Lanes 2-7: Δ*hrpA* expressing wild-type or mutant HrpA on a low copy number plasmid under its endogenous promoter and ribosome binding site. Loading control is FtsZ. (**B**) Distribution of wild-type and mutant HrpA-FLAG in sucrose gradients observed with anti-Flag antibodies. (**C**) Polysome traces from untreated, low-dose (5 μg/ml), or high-dose (500 μg/ml) CAM-treated cells. (**D**) Polysome traces from untreated cells.

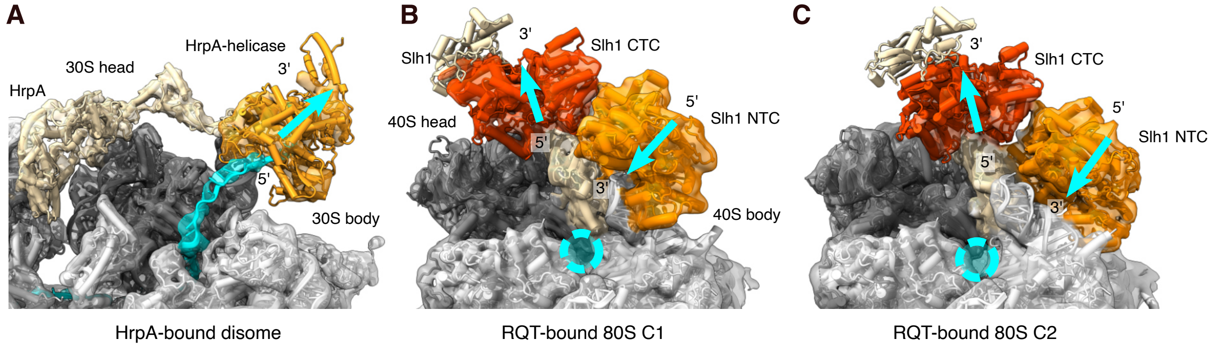

**Fig. S10. Comparison of HrpA-bound disomes with RQT-bound 80S.** (**A**) View of HrpA bound to the *E. coli* disome with emerging mRNA (molecular model fit into the low-pass filtered electron density). The 30S head is shown in dark grey, the 30S in light grey. HrpA is shown in tan except for the helicase domain (orange). The cyan arrow indicates the movement of mRNA during 3’-5’-helicase activity of HrpA. (**B, C**) Same view as **A** showing Slh1 bound to the *Saccharomyces cerevisiae* 80S ribosome[27] in C1 state (POST state with P-site tRNA, **B**) and C2 state (TI-POST-2 state with swiveled 40S head, **C**). The helicase cassettes of Slh1 (NTC; N-terminal cassette and CTC; C-terminal cassette) are shown in orange and red, respectively. The rest of Slh1 is shown in tan. Rqt4 and Cue3 of the yeast RQT complex were omitted for clarity. Cyan arrows indicate the movement of mRNA during 3’-5’-helicase activity of Slh1 and the mRNA entry channel is highlighted by a cyan circle.

### Table S1. Cryo-EM data collection, refinement and validation statistics

| | #1 Class I. (EMD-51318) (PDB 9GFT) | #2 Class II (EMD-51340) (PDB 9GGR) |
|---|---|---|
| **Data collection and processing** | | |
| Magnification | 165000 | 165000 |
| Voltage (kV) | 300 | 300 |
| Electron exposure (e–/Å$^2$) | 40 | 40 |
| Defocus range (μm) | 0.5-2.4 | 0.5-2.4 |
| Pixel size (Å) | 0.727 | 0.727 |
| Symmetry imposed | C1 | C1 |
| Initial particle images (no.) | 2,008,477 | 2,008,477 |
| Final particle images (no.) | 17,182 | 17,805 |
| Map resolution (Å) | 3.1 | 3.2 |
| FSC threshold | 0.143 | 0.143 |
| **Refinement** | | |
| Initial model used (PDB code) | 7qg8, 7qgh, 6zww, 7k00, 8pkl, AlphaFold P43329 | 7qg8, 7qgh, 6zww, 7k00, 8pkl, AlphaFold P43329 |
| Model resolution (Å) | 3.8 | 4.6 |
| FSC threshold | 0.5 | 0.5 |
| Model composition | | |
| Non-hydrogen atoms | 307414 | 309821 |
| Protein residues | 13317 | 13616 |
| Nucleotide | 9463 | 9463 |
| R.m.s. deviations | | |
| Bond lengths (Å) | 0.003 | 0.004 |
| Bond angles (°) | 0.773 | 0.846 |
| Validation | | |
| MolProbity score | 2.19 | 1.83 |
| Clashscore | 7.54 | 8.93 |
| Poor rotamers (%) | 3.70 | 0.11 |
| Ramachandran plot | | |
| Favored (%) | 95.00 | 94.94 |
| Allowed (%) | 4.94 | 5.05 |
| Disallowed (%) | 0.06 | 0.01 |
